# Neural stem cell–derived extracellular vesicles drive early neuroprotective and anti-apoptotic responses in spinal cord injury organotypic slices

**DOI:** 10.64898/2026.05.11.718900

**Authors:** Kristyna Sintakova, Vojtech Sprincl, Ivan Arzhanov, Ruslan Klassen, Lukas Valihrach, Nataliya Romanyuk

**Author notes:** Correspondence: Nataliya Romanuyk.

## Abstract

Spinal cord injury (SCI) is a devastating neurological condition with limited regenerative capacity. Stem cell–based approaches have emerged as promising strategies due to their neuroprotective and immunomodulatory properties, largely mediated by small extracellular vesicles (sEVs) and their molecular cargo, including miRNAs. In this study, we aimed to evaluate the neuroprotective and anti-apoptotic potential of sEVs derived from SPC-01 and iMR-90 neural stem cell sources using an in vitro rat model of SCI. sEVs were isolated from conditioned media and characterized by multi-angle dynamic light scattering and Western blot analysis. Organotypic spinal cord slices (SCS) were used as an *in vitro* SCI model, with injury induced at 18–20 days, followed by immediate sEV application. After 72 h, tissue samples were collected and tissue was analyzed for markers of apoptosis, cytoskeletal integrity, and survival-related signaling pathways. Results show that SCI induced cytoskeletal disruption and increased apoptotic markers. Treatment with sEVs mitigated these changes, reducing injury-associated protein levels toward baseline. Both SPC-01- and iMR-90-derived sEVs exerted comparable neuroprotective effects, accompanied by decreased PTEN expression, enhanced STAT3 phosphorylation, and increased levels of the anti-apoptotic protein Bcl-xL. In parallel, reduced Nogo-A expression and normalization of RhoA suggested improved cytoskeletal stability and attenuation of inhibitory signaling. Together, these findings demonstrate that neural stem cell–derived sEVs promote early neuroprotective responses *in vitro* by modulating key signaling pathways, reducing apoptosis, and stabilizing cytoskeletal dynamics, supporting their potential as a cell-free therapeutic strategy for SCI.

## 1 Introduction

Spinal cord injury (SCI) is a debilitating condition that results in severe neurological impairment, affecting a large number of individuals annually. The impact of SCI includes both temporary and permanent alterations in spinal cord structure, as well as loss of motor, sensory, and autonomic functions (Alizadeh et al., 2019; Dumont et al., 2001). Following a traumatic primary injury, an inflammatory cascade of secondary injury unfolds, encompassing demyelination, ischemia, activation of immune mechanisms, and the release of cytokines and cytotoxic agents. These processes, along with apoptotic and necrotic cell death, contribute to the inflammatory environment at the injury site and promote the spread of damage to the initially healthy tissue surrounding the lesion (Ahuja, Nori, et al., 2017; Hayta & Elden, 2018; Oyinbo, 2011).

Effective treatment for SCI is still unavailable. Major challenges include the fusion of cystic cavities and the formation of a glial scar, the limited regenerative capacity of the central nervous system (CNS), and poor axonal remyelination (Ahuja, Nori, et al., 2017). Current therapeutic interventions are aimed at mitigating SCI consequences, such as cerebrospinal fluid (CSF) drainage to reduce post-injury pressure (Martirosyan et al., 2015), or the use methylprednisolone sodium succinate in combination with surgical decompression (Fehlings et al., 2017). One of the promising experimental therapeutic approaches for SCI is the stem cell transplantation. Mesenchymal stem cells (MSCs) can modulate inflammation at both local and systemic levels in SCI pathology (Quertainmont et al., 2012). Neural stem cell (NSC) transplantation has shown neuroprotective and immunomodulatory effects in CNS pathologies, including SCI, promoting remyelination and functional recovery (Cheng et al., 2016; Martino et al., 2011; Pereira et al., 2019). The positive effect of NSC transplantation in SCI pathology has been confirmed by our previous research (Amemori et al., 2013; Romanyuk et al., 2015). However, persistent challenges with stem cell transplantation include low cell survival rate, tumorigenesis, cell dedifferentiation, and transplant rejection (Balsam et al., 2004; Jeong et al., 2011).

Despite well-documented beneficial effects of stem cells in CNS pathologies, including SCI, their underlying mechanisms remain insufficiently understood. Studies suggest that the neuroprotective and supportive effects of transplanted stem cells may not result from direct replacement of damaged cells, but rather from paracrine signaling mechanisms, potentially mediated by small extracellular vesicles (sEVs) (Baglio et al., 2012; Rong et al., 2019; Zhong et al., 2020), and particularly by small non-coding RNA molecules (ncRNAs) encapsulated within these vesicles (Wiklander et al., 2019). sEVs are membrane-bound nanovesicles, ranging from 30 to 150 nm in size, that facilitate intercellular communication by transporting functional proteins, mRNA, miRNA, lipids, and other molecules. The encapsulated cargo within sEVs is protected from degradation, and sEVs are also capable of crossing the blood-brain barrier. These properties make sEVs promising delivery vehicles for therapeutic molecules into recipient cells and a potential alternative to whole stem cell transplantation, offering several advantages over cell-based therapies, including a reduced risk of immunogenicity, uncontrolled differentiation, and proliferation (Abels & Breakefield, 2016; Elahi et al., 2020; Khan et al., 2023; Wiklander et al., 2019; B. Zhang et al., 2016). For example, intravenous administration of stem cell-derived sEVs has been shown to reduce inflammation and support motor function recovery (Maas et al., 2017). MSC-derived sEVs (MSC-sEVs) have demonstrated anti-inflammatory effects, promoted tissue regeneration by enhancing extracellular matrix remodeling (Gurunathan et al., 2019), supported neurogenesis and functional recovery, and reduced apoptosis (Huang et al., 2020; Kang et al., 2019; W. Liu et al., 2019). However, MSC-sEVs, with those derived from other non-CNS-derived cell types, appear to have a limited capacity to induce key neuroregenerative processes in CNS pathologies, such as axonal remyelination and neurogenesis. In contrast, NSC-derived sEVs (NSC-sEVs) have demonstrated better neurogenic, neuroprotective, and anti-inflammatory effects in the treatment of CNS injuries compared to MSC-sEVs (Rong et al., 2019; Webb et al., 2018; Zhong et al., 2020).

Here, we evaluated and compared the therapeutic potential of sEVs derived from two NSC lines using rat spinal cord slices (SCS) as an *in vitro* model of SCI. We focused on key processes associated with post-injury degeneration and repair, including apoptosis, cytoskeletal integrity, and neural tissue preservation. Our results demonstrate that sEV treatment reduced the expression of apoptosis- and pathology-associated proteins while increasing levels of proteins linked to pro-survival and regenerative responses. These findings support the neuroprotective and anti-apoptotic effects of NSC-derived sEVs and highlight their potential as a cell-free therapeutic strategy for spinal cord injury.

## 2 Materials and Methods

### 2.1 Cell culture

Using well established protocols, two NSC lines were cultivated: human conditionally immortalized spinal fetal cell line (SPC-01) (Pollock et al., 2006) and neural precursors differentiated from human-induced pluripotent (iPSC-NPs) cells derived from fetal lung fibroblast line (iMR-90; ATCC, Manassas, VA, United States) (Polentes et al., 2012). iMR-90 cells were cultivated in T75 or T175 flasks (Nunc, Thermo Fisher Scientific, Roskilde, Denmark) coated with poly-L-ornithine (Sigma-Aldrich, St. Louis, MO, United States); and laminin (20 μg/ml in DMEM:F12) (Sigma-Aldrich, St. Louis, MO, United States); 20 μg/ml in DMEM:F12) in CO2 incubator (MCO-170AICUVH-PE, Panasonic) at 37°C. The medium used for iMR-90 was of following composition: DMEM: F12 and neurobasal medium (1:1), supplements B27 (1:50) and N2 (1:100), a mixture of penicillin/streptomycin antibiotics (1:100) (all from Thermo Fisher Scientific, Waltham, MA, United States), growth factors EGF (10 ng/ml), bFGF (10 ng/ml) and BDNF (10 ng/ml) (all from PeproTech, London, United Kingdom), antibiotics primocin (1:500) (InvivoGen, San Diego, CA, United States). SPC-01 are spinal cord neural precursors, obtained by material transfer agreement in frame of collaboration with The James Black Centre, Department of Neuroscience, King’s College London, 125 Coldharbour Lane, London, UK. The isolation and immortalization of this cell line is described in Pollock et al., 2006. SPC-01 cells were cultivated in T75 and T175 flasks coated with laminin (at the same concentration used in iMR-90 cultivation) in CO2 incubator at 37°C. The culture medium consisted of DMEM:F12 supplemented with human serum albumin (HSA) (0.03%) (Baxter Healthcare Ltd., Norfolk, UK); L-glutamine (2 mM), human apo-transferrin (100 μg/ml), putrescine dihydrochloride (16.2 μg/ml), human insulin (5 μg/ml), progesterone (60 ng/ml), sodium selenite (selenium) (40 ng/ml), EGF (20 ng/ml), 4-hydroxy-tamoxifen (4-OHT) (100 nM) (all from Sigma-Aldrich); bFGF (10 ng/ml) (PeproTech, London, UK), a mixture of penicillin/streptomycin antibiotics (1:100) (InvivoGen, San Diego, CA, United States).Both iMR-90 and SPC-01 had the cultivation medium replaced every other day and the cultures were passaged every 3 days when they reached 70% - 90% confluence. NSC-sEVs used in the experiments were isolated from culture medium of SPC-01 cells at passages 25-30 and from the culture medium of iMR-90 cells at passages 15-20. The medium intended for NSC-sEVs isolation was collected after 72 hours of cultivation. To generate a cellular control for sEV-specific markers in Western blot analysis, SPC-01 and iMR-90 were cultured in 6-well plates (TPP, Trasadingen, Switzerland) under the same conditions as described above.

### 2.2 Vesicle isolation and characterization

Following cultivation, culture media was collected, and sEVs were isolated using sucrose cushion ultracentrifugation, which was evaluated as the best method for sEVs isolation by trial experiments. Cell growth medium was collected and centrifuged at 300 × g for 10 minutes at 4°C, followed by centrifugation at 2000 × g for 10 minutes at 4°C on a 5, 804 R centrifuge (Eppendorf, Hamburg, Germany). The supernatant was transferred to a polycarbonate ultracentrifuge UltraClearTM tube (Beckman Coulter, Inc., Brea, California, United States) and centrifuged for 30 minutes at 4°C and 10 000 × g using Optima XPN-90 Ultracentrifuge (Beckman Coulter, Inc., Brea, California, United States). Subsequently, sEVs were purified through centrifugation with a 30% sucrose cushion for 90 minutes at 4°C and 100 000 × g. The sucrose cushion was composed of 0, 3 g/ml sucrose (Sigma-Aldrich, St. Louis, MO, United States), 24 mg/ml Tris (Roche, Basel, Switzerland), diluted in D2O (Sigma-Aldrich, St. Louis, MO, United States) to final volume of 50 ml, with solution pH adjusted to pH 7,4. Following the centrifugation, the supernatant was carefully aspirated, and the remaining volume (approximately 10 ml) was diluted in phosphate buffered saline (PBS) and centrifuged again at 4°C and 100 000 × g for 90 minutes to remove sucrose from the isolated sEVs. The pellet was then dissolved in 50-100 μl of PBS, and the solution was stored at 4°C. For long-term storage, samples were frozen at − 20 °C.

### 2.3 MADLS

Multi-angle dynamic light scattering (MADLS) measurements were performed using a Zetasizer Ultra instrument (Malvern Panalytical, UK) to determine the size of isolated sEVs, and the ZS XPLORER software was used for data interpretation. For the measurement, a side laser source positioned at a 90° angle, was utilized, and correlogram characteristics were monitored to ensure proper measurement quality. The size of particles presented as median ± interquartile range (IQR).

### 2.4 Western Blot

For Western Blot analysis, cells or tissue samples were homogenized on ice in radio immunoprecipitation assay (RIPA) lysis buffer [150 mM NaCl, 50 mM Tris (pH 8), 1% Triton X-100, 0.5% sodium deoxycholate, 0.1% sodium dodecyl sulfate] (all from Sigma-Aldrich, St. Louis, MO, United States) containing a protease and phosphatase inhibitor (Thermo Fisher Scientific, Waltham, MA, United States). After homogenization, samples were incubated at 4°C for 40 min, followed by a centrifugation at 14,000 RPM and 4°C for 20 min on a 5,804 R centrifuge. Samples were then frozen at − 80°C. To determine total protein concentration in samples, Pierce TM BCA Protein Assay Kit (Thermo Fisher Scientific, Waltham, MA, United States) was utilized according to the manufacturer’s instructions. Spectrophotometric measurements were performed using i-control software on an Infinite® 200 PRO Multimode Reader (Tecan, Mannedorf, Switzerland). NanoDrop 8000 Spectrophotometer (Thermo Scientific) was used for small sample volumes (<50 µl).

Both sEVs and cell samples were diluted in sodium dodecyl sulfate (SDS) sample buffer for denaturation prior to SDS-PAGE analysis [80 mM Tris (pH 6.8), 2% SDS, 10% glycerol, 0.0006% bromophenol blue, 0.1 M DTT] (all from Sigma-Aldrich, St. Louis, MO, United States) and incubated for 5 min at 95°C. Sample volume per well and its protein content were adjusted according to the BCA protein assays results. Electrophoresis was performed using Mini-PROTEAN TGXTM Precast Gels (Bio-Rad, Hercules, CA, United States) with a gradient (8–14%) in electrophoresis buffer (25 mM Tris, 192 mM glycine, 0.1% SDS) (all from Sigma-Aldrich, St. Louis, MO, United States). Each well within the gel was loaded with 10 μg of total protein. The runtime was approximately 30-45 minutes. Following electrophoresis, proteins were then transferred from gels to nitrocellulose membranes (Cytiva, Marlborough, MA, United States) in transfer buffer [25 mM Tris, 192 mM glycine, pH 8.3; along with 20% methanol (v/v)] (all from Sigma-Aldrich, St. Louis, MO, United States). Following transfer, membranes were stained with Ponceau S staining solution (Cell Signaling Technology, Danvers, MA, United States) for protein visualization. Tris-buffered saline / Tween-20 (TBST) buffer (20 mM Tris, 150 nM NaCl, 0.1% Tween 20; pH 7.5) (all from Sigma-Aldrich, St. Louis, MO, United States) was used to wash the membranes between protocol steps. Membranes were incubated with 5% milk solution (Cell Signaling Technology, Danvers, MA, United States) for 1 hour at room temperature in order to block non-specific binding sites. After washing with TBST, membranes were incubated overnight at 4°C with primary antibodies, which are described in **Supplementary table 1.** Primary antibodies were diluted in 5% bovine serum albumin (BSA) solution (Cell Signaling Technology, Danvers, MA, United States), or in 5% milk solution (Cell Signaling Technology, Danvers, MA, United States), or directly in TBST. Following overnight incubation, the membranes were washed with TBST and incubated with the appropriate secondary antibody, diluted in TBST 1:10 000 (**Supplementary table 2**) for 1 hour at room temperature, followed by additional washes with TBST. Super Signal TM West Dura kit (Thermo Fisher Scientific, Waltham, MA, United States) was utilized to visualize the staining. Blots were captured using the Azure c600 (Azure Biosystems, Dublin, CA, United states) in Capture software, and image digitalization and analysis were performed with Fiji software. The results were normalized for endogenous control proteins: vinculin (124 kDa) or β-actin (42 kDa), depending on the molecular weight of the particular protein. At minimum of 3—4 technical replicates were performed for each protein at each experimental time point.

The normalization strategy was as follows: band intensities of target proteins were first normalized to the corresponding endogenous control (β-actin or vinculin). Subsequently, within each experimental group (control, SCI, SCI+sEVs), values were normalized to the control sample, such that the control was set to 1 and all other values are expressed relative to it.

Experiments involving the two types of sEVs were performed separately; therefore, the corresponding bands are presented separately. As no significant differences were observed between SCI samples across experiments, we combined the quantifications to avoid redundant representation of repeated values within each experimental group.

### 2.5 RT-qPCR

SPC-01-sEVs cargo was analyzed using reverse-transcription quantitative polymerase chain reaction (RT-qPCR). Total RNA, including a small fraction of miRNAs, was isolated using the miRNeasy Serum/Plasma Advanced Kit (Qiagen, Hilden, Germany). miRNAs were quantified using two-tailed RT-qPCR assays developed in the Gene Expression Laboratory (Androvic et al., 2017). Following an established protocol, a two-tailed RT-qPCR panel consisting of two synthetic spike-in molecules and three endogenous miRNAs was used for quality control (QC). This QC panel assessed the efficiency of the RNA extraction protocol, the quality of the isolated miRNAs, inhibition rate, and the technical accuracy of the obtained results.

### 2.6 Spinal cord slices as an *in vitro* SCI model

All experiments were designed according to the 3-R rules. The experimental procedures were approved by the Ethics Committee of the IEM ASCR and are in accordance with the Directive of the European Commission of November 24, 1986 (86/609/EEC) on the use of animals in research. Spinal cord slices were used as an *in vitro* model of SCI. Slices were prepared from 5-7-day old male Wistar rats. After administering the anesthetic Vetbutal (Virbac, Carros, France), the animals were euthanized by decapitation, and their spinal cords were extracted. The isolated spinal cord tissue was placed on 4% agar (Sigma-Aldrich, St. Louis, MO, United States) blocks on a Leica VT1200s vibratome (Leica Microsystems, Wetzlar, Germany), positioned along the blade. The agar, vibratome, and PBS wash were kept on ice. The spinal cord was sliced into 1 cm longitudinal sections, each 350 µm thick. These sections were placed on membrane inserts (MilliporeSigma-Aldrich, Burlington, MA, United States) in a 6-well plate (TPP, Trasadingen, Switzerland), with three to four slices on each insert. SCS culture media [DMEM/F12, HBSS (both from Gibco, Grand Island, New York, United States), Horse serum (EastPort LifeSciences, Prague, Czech Republic), HEPES (Sigma-Aldrich, St. Louis, MO, United States), L-glutamine, glucose, amphotericin B, and mixture of penicillin/streptomycin antibiotics (1:100) (all from Thermo Fisher Scientific, Waltham, MA, United States] was added. SCS were then cultivated under humidified conditions with 5% CO2 at 34 °C for 2-3 weeks.

### 2.7 Alamar Blue assay

To assess the viability of SCS and determine the most suitable and reliable time period for an *in vitro* experiment, Alamar Blue assay was performed every 3 days for 3 weeks after extraction. Slices were cultured in 10% Alamar Blue (7-Hydroxy-3H-phenoxazin-3-one-10-oxide sodium salt) (Sigma-Aldrich, St. Louis, MO, United States) in SCS medium. Following 3 hours of incubation, fluorescence level of reduced Alamar Blue was measured with Infinite® 200 PRO Multimode Reader and *i-control* software (Tecan, Männedorf Switzerland) with an excitation wavelength of 550 nm and an emission wavelength of 590 nm (Petrenko et al., 2005). The ratio between the fluorescence of experimental and blank samples (the same medium without cells) was used as the Alamar Blue value. Data were presented as relative fluorescence units (RFU).

### 2.8 Immunohistochemistry

For analysis of the tissue structure of the slices, immunohistochemical staining was performed. The slices were fixed with 4% paraformaldehyde (Penta, Prague, Czech Republic) and washed with PBS (Thermo Fisher Scientific, Waltham, MA, United States), followed by permeabilization with 0.5% Triton X-100 (Sigma-Aldrich, St. Louis, MO, United States) diluted in PBS for 20 minutes in the dark at room temperature, and then washed in 0.2% Triton X-100 in PBS. Non-specific staining was blocked by incubating the slices in a solution of 0.3M glycine (Sigma-Aldrich, St. Louis, MO, United States), 10% NGS, and 0.2% Triton X-100 in PBS for 2 hours at room temperature, protected from light. After washing with PBS, slices were incubated overnight at 4°C in the dark with primary antibodies diluted in 10% NGS, 0.2% Triton X-100 in PBS. Individual antibodies and their dilutions are described in **Supplementary table 1**. Following incubation, slices were washed with PBS, then in 0.2% Triton X-100 in PBS solution for 10 minutes, and incubated with secondary antibodies (diluted 1:400 in the same solution as the primary antibodies, (**Supplementary table 2**) for 2 hours at 4°C. For cell nuclei staining, 4′,6-diamidino-2-phenylindole (DAPI) (Sigma-Aldrich, St. Louis, MO, United states) (1:2500) in 0.2% Triton X-100 in PBS was applied. The slices were then carefully removed from the plate wells, mounted onto microscopic coverslips using Aqua Poly/Mount (Polysciences, Warrington, PA, United States), and covered with cover glasses. Immunocytochemical staining was captured using a Carl Zeiss LSM 880 NLO confocal microscope (Carl Zeiss AG, Oberkochen, Germany), equipped with Argon/Helium/Neon lasers. Markers specific to spinal cord cells were detected, confirming the preservation of neurons, astrocytes, and microglia in the slices.

### 2.9 SCI *in vitro* model and sEV treatment

Based on the results of the Alamar Blue assay, experiments evaluating the effect of sEVs in SCS with induced SCI were conducted between days 18 and 21 after SCS preparation. The slices were divided into three groups: (1) a control group without induced SCI and without sEV treatment; (2) an SCI group, and (3) a group with induced SCI and treated with sEVs. Incomplete SCI was induced using spring scissors; (Fine Science Tools, Foster City, CA, United States) approximately at the center of the spinal cord cross-section. The dose of sEVs for SCI treatment was calculated based on total protein content at 50 µg/ml of culture medium. sEVs samples were diluted in filtered SCS culture medium and the medium was changed immediately after lesion induction. After 3 days of cultivation, the slices were collected for Western Blot analysis, following the same protocol as described above. Protein concentration in samples determined using the BCA protein assay. Western Blot was employed to assess changes in the expression levels of apoptotic and neuron-specific markers, as well as to evaluate the anti-apoptotic and neuroprotective effects of sEVs on neural tissue following SCI. The same protocol as used for sEVs analysis was applied, using of PVDF membranes (Thermo Fisher Scientific, Waltham, MA, United States) instead of nitrocellulose membranes.

### 2.10 Statistical analysis

Western Blot data were normalized to the corresponding endogenous protein control. Samples were analyzed in groups of four, with the control sample set to 1 and the spinal cord injury and sEVs-treated samples normalized relative to the control. Data from WB quantification, as well as metabolic activity data are shown as the mean ± STD from three or more independent experiments. Statistical analysis of the differences between groups was performed using repeated measures ANOVA. Significant difference between compared groups were considered when p<0.05 *; p<0.01 **, p<0.001 ***, p<0.0001 ****.

## 3 Results

### 3.1 Characterization of sEVs isolated from SPC-01 and iMR-90 culture media

sEVs were isolated from conditioned media collected every 3 days from SPC-01 and iMR-90 cell cultures using ultracentrifugation with a sucrose cushion, which was identified in preliminary experiments as the most efficient isolation method. To confirm their identity as sEVs, isolated particles were analyzed by MADLS and Western Blot analysis. MADLS revealed no significant differences between SPC-01- and iMR-90-sEVs with median diameters of 84.5 ± 14.2 nm and 72.8 ± 13.2 nm, respectively, as shown in **Figure 1A**. Western blot analysis confirmed the presence of exosomal markers Alix, TSG101, CD9, CD63, and CD81 in both sEVs populations, while calnexin, endoplasmic reticulum (ER) marker, was detected only in cell lysates **(Fig. 1B**). Because the SPC-01 cells were immortalized using a *c-myc* gene encoding vector, the presence of the protein product of this gene in SPC-01-sEVs-was assessed to rule out potential oncogenicity but not detected (**Fig. 1C**). RT-qPCR analysis demonstrated the presence of miRNAs within SPC-01-sEVs, including miR-21a-5p and other miRNAs implicated in central nervous system injury-related regulatory processes (miR-20a-5p, miR-24-3p, miR-320a-3p). (**Fig. 1D)**.

**Figure 1.**
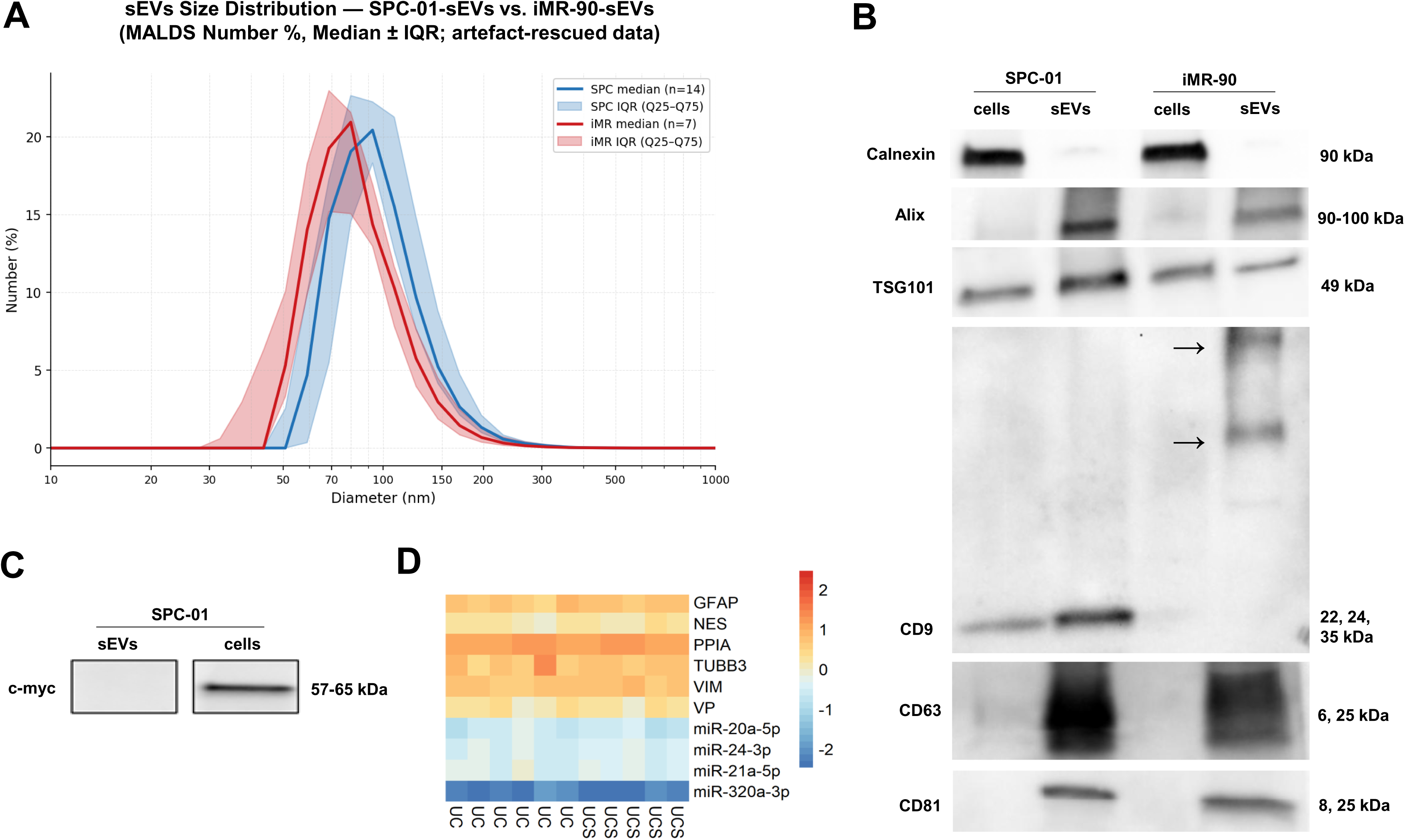
Characterization of sEVs derived from SPC-01 and iMR-90 culture media. Size distribution of isolated particles was assessed by MADLS **(A)**, showing no significant differences in size between SPC-01-sEVs (84.5 ± 14.2 nm) and iMR-90-sEVs (72.8 ± 13.2 nm). Western blot analysis **(B)** confirmed the presence of exosomal markers (Alix, TSG101, CD9, CD63, CD81) in both sEV preparations, while the cellular marker calnexin was detected only in cell lysates. CD9 dimerization (denoted by arrows) was observed in iMR-90-sEVs and CD63 glycosylation was observed in both sEVs samples. The stem cell marker c-Myc was absent in SPC-01-sEVs **(C).** RT-qPCR analysis **(D)** confirmed the presence of selected miRNAs in SPC-01-sEVs. *MADLS, multi-angle dynamic light scattering; TSG101, Tumor Susceptibility Gene 101; CD, cluster of differentiation; GFAP, glial fibrillary acidic protein; DAPI, 4′,6-diamidino-2-phenylindole*; *UC, ultracentrifugation; UCS, ultracentrifugation with sucrose cushion; IQR, Interquartile Range*.

### 3.2 Spinal cord slices retain structural integrity and cellular composition suitable for evaluating sEVs effects after spinal cord injury

To investigate the therapeutic potential of SPC-01- and iMR-90-sEVs and their effects on processes associated with SCI pathology, rat SCI *in vitro* model of organotypic spinal cord tissue slices (SCS) **(Fig. 2A)**. This model is less demanding than *in vivo* SCI models while offering advantages over *in vitro* cell-based models, such as scratch or laser dissection models, as it partially preserves tissue connectivity and extracellular matrix interactions. In addition, its use is consistent with the Three Rs concept (reduction, replacement, and refinement). Prior to the SCI experiment, we characterized the SCS in terms of viability, and structural integrity of the slices was assessed to determine the most reliable time window for *in vitro* experimentation. SCS viability was assessed using the Alamar Blue assay every 3 days over a 3-week period **(Fig. 2B)**. An initial decrease in metabolic activity was observed during the first days after preparation, which was attributed to the elimination of damaged cells generated during slice preparation. Subsequently, metabolic activity gradually recovered and stabilized over the subsequent 10–12 days. This period was defined as the experimental window, representing the most suitable timeframe for *in vitro* experiments. Based on these findings, all subsequent SCI experiments were performed between days 18 and 20 after SCS preparation. Immunohistochemical staining confirmed that major spinal cord cell types were well preserved within the cultured slices, as shown in **Figure 2C**. Astrocytes were identified using glial fibrillary acidic protein (GFAP), S100 calcium-binding protein B (S100β), and vimentin. Neuronal populations were detected using βIII-tubulin, a neuron-specific microtubular protein and neurofilament-H (NF-H), a component of the neuronal intermediate filament cytoskeleton. Changes in microglial and neuronal populations were monitored over the cultivation period (1, 2, and 3 weeks) via Iba1 and NF-H staining, respectively. Staining for Iba1 suggested a reduction in immunocompetent microglial cells in the tissue culture over the course of the observation period **(Fig. 2D)**. Cell nuclei were stained with DAPI. These findings indicate that SCS retain spinal cord tissue–specific morphology and cellular composition during prolonged cultivation and may serve as a suitable *in vitro* model for studying spinal cord injury.

**Figure 2.**
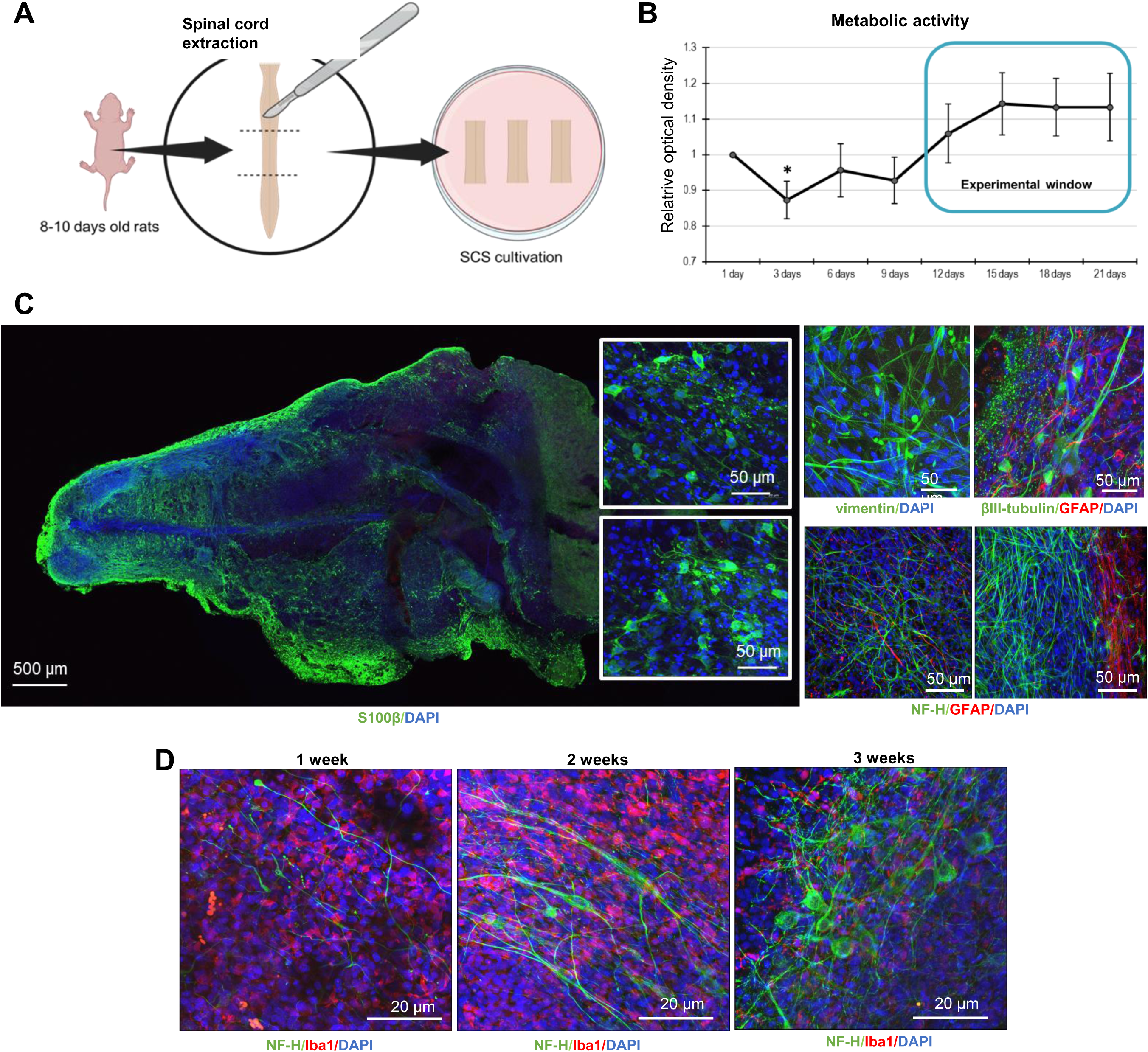
Spinal cord slices and their use as an *in vitro* model of spinal cord injury. Schematic overview of spinal cord slice (SCS) preparation **(A)**. Tissue viability was assessed using the Alamar Blue assay over 3 weeks **(B)**, showing an initial decrease in metabolic activity followed by recovery and stabilization between days 10-12; experiments were therefore performed on days 18–20. Immunohistochemical analysis **(C)** confirmed the preservation of major spinal cord cell types, with astrocytes (GFAP, S100β, vimentin) and neurons (βIII-tubulin, NF-H) detected after 3 weeks in culture. Microglial activity was assessed by Iba1 staining over time (**D**), with a progressive decrease in signal intensity accompanied by improved organization of NF-H-positive axonal structures. *NF-H, Neurofilament-H; GFAP, glial fibrillary acidic protein; S100β, S100 calcium-binding protein B; Iba1, ionized calcium-binding adapter molecule 1; DAPI, 4′,6-diamidino-2-phenylindole*.

### 3.3 sEV application mitigates SCI-induced changes in neurons and astrocytes

To investigate the neuroprotective and anti-apoptotic potential of SPC-01-sEVs and iMR-90-sEVs, we established three experimental groups: control SCS without induced SCI, SCS with incomplete SCI, and SCS with incomplete SCI treated with either SPC-01-sEVs or iMR-90-sEVs. SCI was induced on days 18–20 of cultivation, and slices were harvested 72 hours post-injury for protein analysis. Western blot was performed to evaluate changes in key proteins involved in cytoskeletal organization, apoptotic pathways, and injury-related signaling mechanisms, providing insight into the ability of sEV treatment to counteract injury-induced cellular changes.

Following SCI, the integrity of neuronal and axonal cytoskeletal structures is compromised, resulting in disrupted axonal transport, structural destabilization, and reduced neurite outgrowth. Astrocytes respond with reactive gliosis, contributing to glial scar formation. To assess whether sEV treatment modulates these injury-induced changes, we performed Western blot analysis of key markers of neuronal and axonal structure (MAP2, NF-H), axonal growth regulation (RhoA, Nogo-A), and astrocyte-associated protein S100β, as seen in **Figure 3**. MAP2, a neuron-specific cytoskeletal protein expressed in multiple isoforms, localizes primarily to neuronal cell bodies and dendrites and plays a critical role in microtubule organization (Chung et al., 1996). Here, only the lower-molecular-weight isoform (∼75–82 kDa) was consistently detectable **(Fig. 3A)**, possibly reflecting degradation of the higher-molecular-weight isoform (∼280 kDa) in SCS during prolonged cultivation. The original blot images are provided in Supplementary File 1, and the corresponding raw quantification data are included in Supplementary File 2. NF-H, an axonal cytoskeletal protein, provides structural support for neurons (Al-Chalabi & Miller, 2003), and its elevation in CSF is a marker of post-SCI axonal damage and neurofilament breakdown (Petzold, 2005). **Figure 3A** and **3B** show that MAP2 and NF-H were significantly increased following SCI, while sEV treatment resulted in significant reduction of both markers. Next, astrocytic marker S100β was examined to evaluate astrocyte response, as it regulates intracellular and extracellular calcium homeostasis, with elevated levels indicating reactive astrogliosis and glial scar formation. As shown in **Figure 3C**, S100β levels were slightly decreased in the SCI group, but increased following sEV treatment, with a significant upregulation observed in the iMR-90-sEVs group. RhoA, a small GTPase regulating actin dynamics and cellular shape, was due evaluated to its role in growth cone collapse and inhibition of neurite outgrowth via the RhoA/ROCK pathway (Fujita & Yamashita, 2014). Similarly, Nogo-A, a high-molecular-weight myelin-associated protein expressed in oligodendrocytes and neurons, restricts axonal growth and neurite outgrowth via the same pathway (Wälchli et al., 2013). The results show Nogo-A levels were elevated following SCI and returned to control levels in the sEV-treated groups **(Fig. 3D)**. In contrast, RhoA was significantly decreased in the SCI group but increased following sEV treatment, with a significant elevation observed in the SPC-01-sEVs treated group **(Fig. 3E)**. These results suggest that NSC-sEVs modulate neuronal and glial responses after injury, contributing to the preservation of cytoskeletal integrity.

**Figure 3.**
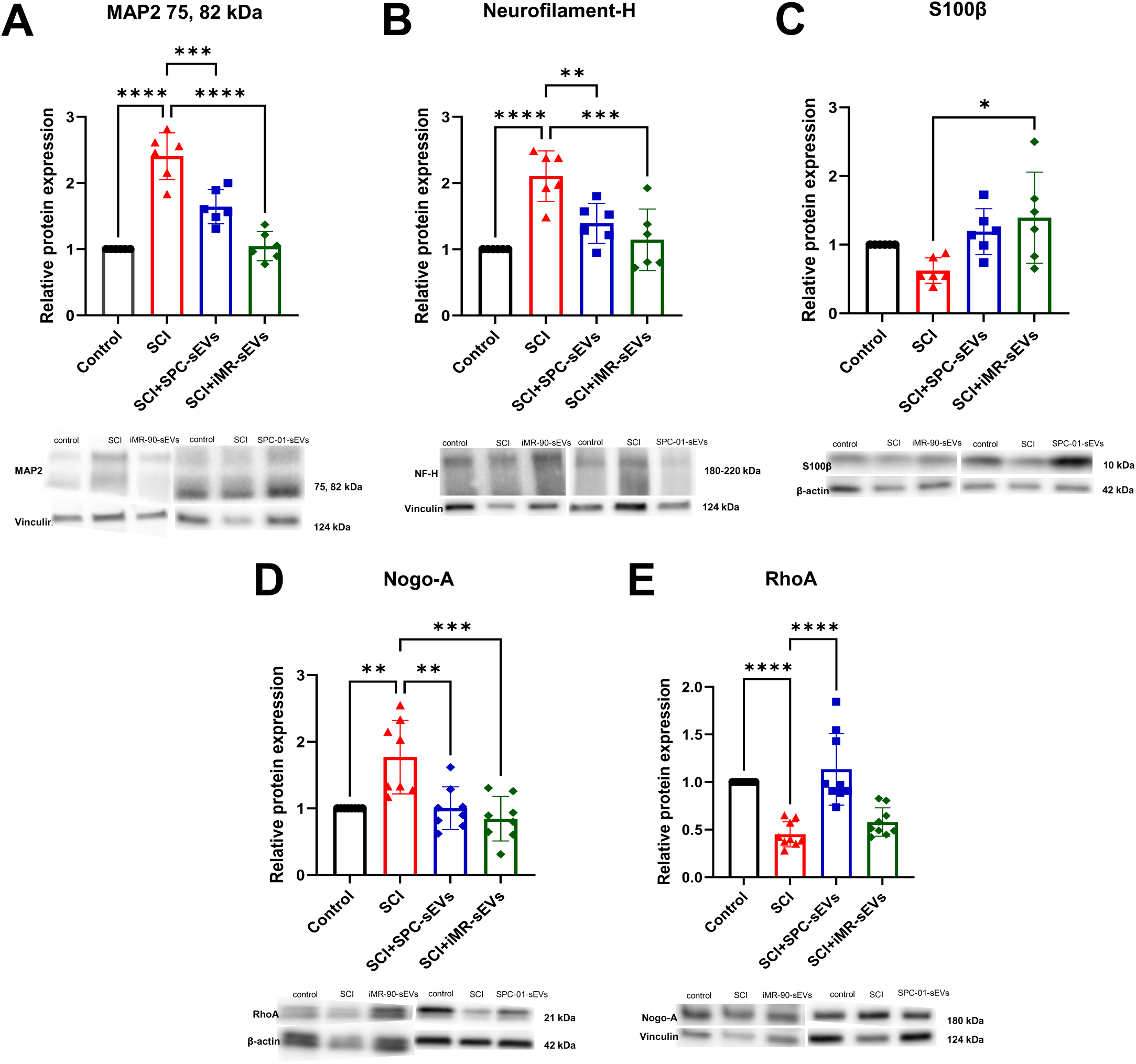
Modulation of cytoskeletal and glial proteins in spinal cord slices following SCI and sEVs treatment. Spinal cord slices (SCS) were divided into three groups: control, SCI, and SCI treated with SPC-01-sEVs or iMR-90-sEVs. After 72 h, slices were harvested for Western Blot analysis. SCI significantly increased, MAP2 **(A)**, NF-H **(B)**, and Nogo-A levels **(D)**. S100β was slightly decreased **(C)** and RhoA was markedly reduced **(E)**. Treatment with sEVs decreased NF-H and MAP2, restored Nogo-A to control levels, and increased RhoA and S100β. Protein levels were normalized to vinculin or β-actin. All data represent mean ± STD. The level of statistical significance was marked as follows: p<0.05 *; p<0.01 **, p<0.001 ***, p<0.0001 **** (n=3-6). *MAP2, Microtubule-associated protein 2; NF-H, Neurofilament-H; S100β, S100 calcium-binding protein B; RhoA, Ras homolog family member A; Nogo A, Neurite Outgrowth inhibitor-A*.

### 3.4 sEVs reduce apoptosis and stabilize mitochondrial pathways after SCI

Apoptotic cell death following SCI occurs in glial cells, neurons, and oligodendrocytes at and beyond the injury site, starting within hours and persisting for weeks. This process involves both intrinsic (mitochondrial) and extrinsic (death receptor) pathways, ultimately leading to caspase activation and DNA fragmentation. The intrinsic pathway is regulated by the balance between pro-apoptotic Bcl-2-associated X protein (Bax) and anti-apoptotic B-cell lymphoma-extra-large (Bcl-xL). Bcl-xL inhibits Bax, thereby preserving mitochondrial integrity and ATP production under stress, whereas Bax promotes cytochrome C release, leading to activation of caspase-9 and downstream executioner caspase-3(Abbaszadeh et al., 2020). Caspase-3 is synthesized as an inactive proenzyme (∼35 kDa) and becomes activated through cleavage into subunits of ∼17 and 19 kDa (Casili et al., 2020). Following SCI, cleaved caspase 3 and Bax levels were markedly increased (**Fig. 4A, B),** while Bcl-xL was decreased **(Fig. 4C)**. Administration of sEVs reduced cleaved caspase-3 and Bax levels in both treatment groups; however, an increase in Bcl-xL expression was observed only in the SPC-01-sEV-treated group. These findings indicate a reduction in apoptotic signaling following sEV treatment.

**Figure 4.**
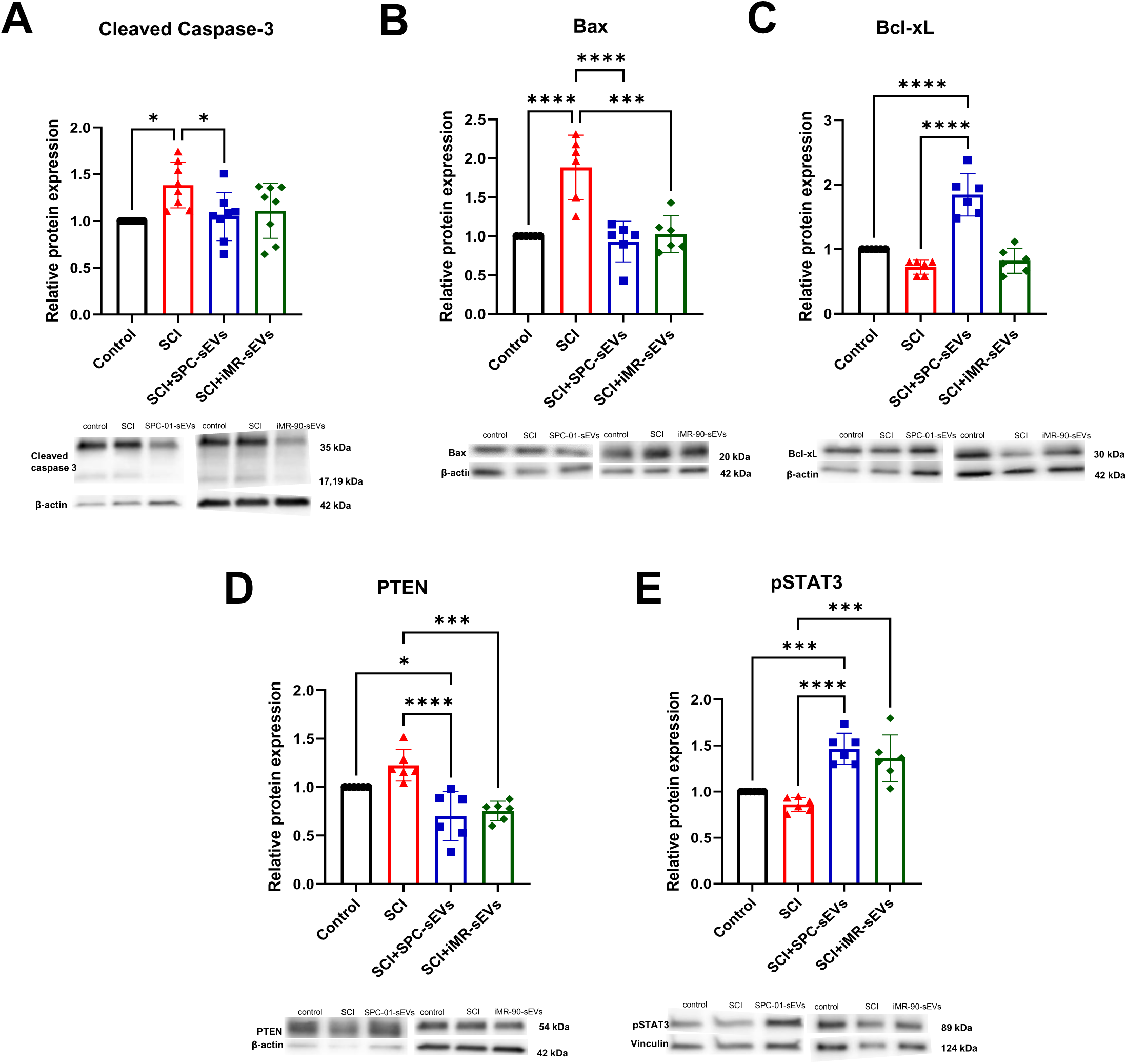
Regulation of apoptotic and survival signaling following sEVs treatment after SCI. PTEN, a negative regulator of the PI3K/Akt/mTOR pathway, was increased after SCI **(D)**, together with the apoptotic marker cleaved caspase-3 **(A)** and decreased following sEV treatment. Expression of proteins involved in apoptosis (Bax and Bcl-xL) shifted toward an anti-apoptotic profile after treatment **(B, C)**. pSTAT3 was significantly elevated following sEV administration **(E)**. Overall, these changes indicate suppression of apoptosis and activation of survival signaling. Protein levels were normalized to vinculin or β-actin. All data represent mean ± STD. The level of statistical significance was marked as follows: p<0.05 *; p<0.01 **, p<0.001 ***, p<0.0001 **** (n=3-6). *Bax, Bcl-2-associated X protein; Bcl-xL, B-cell lymphoma-extra-large; PTEN, phosphatase and tensin homolog; pSTAT3, phosphorylated signal transducer and activator of transcription 3*.

### 3.5 sEVs activate neuroprotective and regenerative signaling pathways after SCI

Key signaling pathways, including PI3K/Akt/mTOR and JAK2/STAT3, are critical for the cellular response to SCI. PI3K/Akt activation protects against oxidative stress, inflammation, and apoptosis, while promoting neurite regeneration and neuronal survival. Phosphatase and tensin homolog deleted on chromosome 10 (PTEN) is a tumor suppressor gene and a negative regulator of PI3K/AKT pathway. Inhibition of PTEN following SCI can activate PI3K/Akt signaling, promoting glial and neuronal responses that support tissue repair, and PTEN deletion enhances axonal regrowth and neuronal survival (C. L. Xiao et al., 2022). Here, PTEN levels were slightly increased in SCI group. Following sEV treatment, PTEN levels were markedly reduced in both groups **(Fig. 4D)**.

Signal transducer and activator of transcription 3 (STAT3) is a transcription factor expressed in neurons, astrocytes, and other CNS cell types. Upon phosphorylation by Janus kinases (JAK), phosphorylated STAT3 (pSTAT3) dimerizes and translocates to the nucleus to promote neuronal survival and axonal growth. In astrocytes, acute STAT3 activation regulates reactive gliosis, scar formation, and limits inflammation after SCI (Herrmann et al., 2008). On Western blot, STAT3 appears as STAT3α (∼86 kDa) and STAT3β (∼79–80 kDa); the ∼86 kDa isoform was analyzed in this study **(Fig. 4E)**. No significant change in pSTAT3 was observed in SCI slices. sEV-treated groups exhibited significant upregulation, consistent with activation of protective and regenerative signaling pathways. Our results show that there we no significant changes in pSTAT3 levels in SCI group. SPC-01- and iMR-90-sEVs treated groups exhibited a significant upregulation of pSTAT3, indicating engagement of survival and regenerative signaling.

Taken together, these results indicate that sEV treatment modulates the acute SCI environment, as reflected by changes in cytoskeletal markers, downregulated pro-apoptotic signaling, and enhanced protective and survival pathways. SPC-01-sEVs produced a stronger effect across multiple markers, suggesting greater neuroprotective potential. Overall, the findings highlight the potential of sEVs to stabilize injured spinal cord tissue and mitigate early injury-induced disruptions.

## 4 Discussion

Spinal cord injury is a severe neurological condition with limited therapeutic options, as current approaches treatments primarily target secondary damage rather than restore lost neural function. Stem cell-based therapies have shown promise due to their immunomodulatory, anti-apoptotic, and neuroprotective properties, much of which appears to be mediated by paracrine signaling via small extracellular vesicles. While MSC-derived sEVs retain the anti-inflammatory and pro-angiogenic properties of their parent cells, along with other non-CNS cell exhibit limited capacity to support neuroregeneration in SCI, including remyelination, axonal regeneration, neurogenesis, and neuronal plasticity. In contrast, CNS-derived sEVs demonstrate neuroprotective and pro-regenerative effects in experimental CNS injury models, promoting functional recovery, plasticity, neurogenesis, and suppressing neuroinflammation (Amemori et al., 2013; Cheng et al., 2016; Dutta et al., 2021; Romanyuk et al., 2015). Few studies have directly compared the effects of sEVs derived from different cell sources in SCI. Most first evaluate whole-cell transplantation (e.g., umbilical, mesenchymal, or neural stem cells), followed by sEVs derived from these cells, sometimes enriched with neuroprotective miRNAs such as miR-133, a PTEN inhibitor (D. Li et al., 2018), or administered via different routes (Kostennikov et al., 2022). Comparative analyses of sEVs from multiple sources have been reported in other contexts (Pishavar et al., 2021; K. Zhang & Cheng, 2025), and our group previously examined hAT-MSC, SPC-01, and iMR-90 transplantation in SCI *in vivo* (Ruzicka et al., 2017). Building on this, the present study evaluates NSC-derived sEVs from SPC-01 and iMR-90 in a rat SCS *in vitro* model of SCI, assessing their effects on cytoskeletal integrity, apoptotic signaling, and key neuroprotective pathways, including PI3K/Akt/PTEN and STAT3.

sEVs are nanosized membrane-bound vesicles (30–150 nm) released by most cell types that mediate intercellular communication through the transfer of proteins, lipids, and nucleic acids, influencing recipient cell responses under both physiological and pathological conditions (Théry et al., 2018). In this study, sEVs were isolated by ultracentrifugation with a sucrose cushion, yielding consistent particle size, concentration, protein content, and characteristic exosomal markers. MADLS analysis **(Fig. 1A)** confirmed that most particles fell within the size range defined by MISEV guidelines (Welsh et al., 2024), with no significant differences between SPC-01-sEVs (84.5 ± 14.2 nm) and iMR-90-sEVs (72.8 ± 13.2 nm). Western blot detected established exosomal markers Alix, TSG101, CD9, CD63, and CD81 in both vesicle types **(Fig. 1B)**. These proteins are commonly associated with sEV biogenesis and function. CD9, CD63, and CD81 belong to the tetraspanin family, while Alix and TSG101 are components of the endosomal sorting complex required for transport (ESCRT) (Colombo et al., 2014). In contrast, the ER marker calnexin was detected only in cell lysates and not in vesicle samples, indicating minimal contamination by intracellular components. Heavy glycosylation of CD63 was observed, consistent with known post-translational modifications in sEVs (Kowal et al., 2017). CD9 in iMR-90-sEVs appeared at a higher molecular weight than in SPC-01-sEVs, likely due to post-translational modifications or protein complex formation commonly observed for tetraspanins. *c-Myc*, a proto-oncogene used to immortalize SPC-01 cells *(Pollock et al., 2006),* was not detected in SPC-01-sEVs **(Fig. 1C)**, alleviating concerns of oncogenic transfer. The neuroprotective and immunomodulatory properties of sEVs are mediated in part by small non-coding RNAs, particularly miRNAs, which have been implicated in multiple aspects of SCI pathology and recovery *(Nieto-Diaz et al., 2014)*. Previous studies examining stem cell-derived vesicles have identified miRNAs associated with neuroprotection and regeneration (Chen et al., 2021; Kang et al., 2019; Z. Wang et al., 2020). In the present study, RT-qPCR analysis revealed miRNAs in SPC-01-sEVs, including miR-20a-5p, miR-320a-3p, miR-24-3p, and miR-21a-5p, which have been associated with regulatory processes in the central nervous system **(Fig. 1D)**. A detailed transcriptomic and proteomic study focusing on both vesicles and producer cells is ongoing (Sprincl et al., in preparation). Taken together, these results confirm successful isolation of functional sEVs populations from both cell types.

To investigate the effects of sEVs on processes associated with SCI, including apoptosis, neuronal survival, and axonal remodeling, SPC-01- and iMR-90-sEVs were applied to rat SCS *in vitro* SCI model (**Fig. 2A)**. SCS preserve key features of native tissue architecture while allowing controlled experimental manipulation. Alamar Blue assay confirmed stable metabolic activity during long-term cultivation, with an initial transient decrease likely reflecting the elimination of cells damaged during preparation, followed by stabilization over 10–12 days, defining a reliable experimental window **(Fig. 2B)**. Experiments were therefore performed between days 18 and 20, when slices displayed metabolic activity. Immunohistochemistry confirmed that SCS retained spinal cord tissue structure throughout the 3-week cultivation period **(Fig. 2C)**. Neurons and astrocytes were preserved, as indicated by βIII-tubulin, neurofilament H, GFAP, and S100β staining. Changes in glial and neuronal populations were monitored using Iba1 to detect microglia and neurofilament H to visualize neurons **(Fig. 2D)**. Iba1, a microglia-specific protein whose expression rises during activation, is widely used as a marker of neuroinflammatory responses after CNS injury, including SCI (Xvlei Hu et al., 2022). Over time, microglial staining gradually decreased, reflecting progressive microglial loss in the absence of systemic immune cell replenishment. These observations should be interpreted in the context of NSC-sEV experiments, taking into account the limitations of the SCS *in vitro* SCI model. Post-SCI pathophysiological responses evolve through acute, subacute, and chronic phases (Ahuja, Wilson, et al., 2017). SCS were harvested for Western blot analysis 3 days post-SCI induction, capturing late acute to early subacute events rather than long-term remodeling processes characteristic of later stages of SCI. By contrast, many *in vivo* studies evaluate therapeutic effects at later time points, often around 28 days (Jia et al., 2021; Rong et al., 2019) or up to eight weeks after injury (Xinyuan Hu et al., 2022), often including functional assessments such as the Basso-Beattie-Bresnahan locomotor scale. Additionally, this *in vitro* SCI model lacks systemic immune interactions and circulating immune cells, consistent with the gradual decline in microglial staining over time. Despite these limitations, the preserved cellular architecture and viability of SCS provide a suitable platform for studying early molecular responses to SCI and the modulatory effects of sEVs on neuronal survival and cytoskeletal integrity. These early molecular responses are considered critical initiating events that can influence later neuronal survival and regenerative capacity, although they do not directly reflect long-term functional outcomes.

Following SCI, neuronal and axonal cytoskeletal integrity is disrupted, resulting in impaired axonal transport, structural destabilization, and reduced neurite outgrowth. This damage is further exacerbated by reactive oxygen species (ROS), which induce oxidative stress, protein misfolding, and fragmentation (Yu et al., 2023). In this study, we examined changes post-SCI induction in neuronal and astrocytic cytoskeletal markers (MAP2, NF-H, S100β) and inhibitory proteins Nogo-A and RhoA to evaluate the effects of SPC-01- and iMR-90-sEVs on structural stability and early neuroprotective responses.

MAP2, a dendrite-specific cytoskeletal protein, is altered following SCI, reflecting dendritic injury and cytoskeletal disruption. Rodent models show rapid MAP2 loss within hours post-contusion, with surviving dendrites often displaying beading patterns indicative of early regenerative responses (S. X. Zhang et al., 2000). In our SCI model, MAP2 levels were markedly elevated 3 days post-SCI **(Fig. 3A)**, likely due to pathological accumulation of damaged protein from necrotic and apoptotic cell death in the late acute to early subacute phase, consistent with the known vulnerability of MAP2 to calpain-mediated degradation which contributes to microtubule destabilization in SCI (DeGiosio et al., 2022; Springer et al., 1997). sEVs treatment restored MAP2 to levels, suggesting microtubule stabilization and normalization of axonal transport. NF-H, a major cytoskeletal protein in large myelinated axons, showed similar patterns: elevated post-SCI, then reduced toward baseline with sEV treatment, indicating preservation of axonal integrity **(Fig. 3B)**. Phosphorylated NF-H (pNF-H) is released into circulation following neuronal injury and serves as a marker of cumulative axonal damage in SCI (Kuhle et al., 2015; Stukas et al., 2023). In SCI patients, pNF-H becomes detectable 12 h post-injury and remains elevated for up to 21 days, unlike the rapid decline of S100β levels (Hayakawa et al., 2012; Kwon et al., 2010). Similar kinetics are observed in rodent models, where NF-H peaks around 72 h post-injury (Rabon et al., 2025), corresponding to the late acute to early subacute phase captured in our SCS model.

Astrocytic marker S100β was analyzed to assess glial responses following SCI. In SCI, S100β elevation is typically transient, peaking within hours post-injury and returning to baseline within 24 h (Haddadi et al., 2024), which may explain the absence of significant expression changes in our model at 3 days post-SCI. Consistently, severe excitotoxic injury induces sharp S100β elevation, whereas mild insults do not, reflecting functional astrocyte stress rather than solely glial number (Mazzone & Nistri, 2014). In line with this, untreated SCI slices in our study exhibited slight decrease in S100β **(Fig. 3C)**, possibly due to astrocyte loss at the lesion site and reduced supportive activity during the acute phase, consistent with the cytoskeletal disruption observed in MAP2 and NF-H **(Fig. 3A, B)**. In contrast, sEVs treatment increased S100β levels, which together with elevated pSTAT3 **(Fig. 4E)** suggests reactivation of astrocytic support and engagement of pro-survival signaling pathways. Previously, MSC-derived sEVs have been shown to modulate astrocytic responses n and promote anti-apoptotic signaling via the miR-21/JAK2/STAT3 pathway (Yang et al., 2024),while neuron-derived exosomes enriched in miR-124-3p suppress activation of pro-inflammatory astrocytic states via PI3K/Akt/NF-κB and STAT3 signaling (Jiang et al., 2020). Overall, our results suggest that SPC-01- and iMR-90-sEVs modulate astrocyte activation toward a state associated with enhanced neutrophic and pro-survival signaling. Through pSTAT3-associated pathways, sEVs may contribute to a supportive glial environment that limits secondary injury processes and supports cytoskeletal stability during the early phase after SCI.

Nogo-A, a myelin-associated protein, inhibits neurite outgrowth and restricts structural plasticity and axonal regeneration and is therefore a relevant therapeutic target in SCI (Schweigreiter & Bandtlow, 2006). In rodent SCI models, its expression is dynamically regulated, peaking around day 7 post-injury before gradually declining by day 14 (J. W. Wang et al., 2015). In our *in vitro* SCI model, Nogo-A levels were already elevated 3 days post-SCI, suggesting an intrinsic neuronal or myelin-derived response even in the absence of systemic immune signaling **(Fig. 3D)**. Treatment with SPC-01- and iMR-90-sEVs restored Nogo-A levels to baseline, indicating attenuation of early inhibitory signaling. Nogo-A exerts its inhibitory effects partly via the RhoA/Rho-associated kinase (ROCK) pathway, a key regulator of cytoskeletal dynamics and axonal growth (Dergham et al., 2002). This pathway is well established as a central mediator of the inhibitory post-injury environment, driven by myelin-associated factors and extracellular matrix components such as CSPGs (Roy et al., 2021) and forms part of the broader secondary injury cascade (Kimura et al., 2021). RhoA activation occurs rapidly post-SCI *in vivo* (Theis et al., 2017) and can persist for days to weeks across multiple neural cell types (Wu & Xu, 2016). However, its regulation is not strictly monotonic. Early time-course studies demonstrate that RhoA expression may transiently decrease shortly after injury before increasing at later stages, with a slight reduction in mRNA levels reported as early as 1 hour post-injury followed by a delayed peak (Sung et al., 2003). In addition, RhoA activity does not necessarily correlate with total protein levels, as substantial increases in active RhoA (RhoA-GTP) can occur without changes in overall expression (Dubreuil et al., 2003). Importantly, these observations highlight that total RhoA levels may not directly reflect pathway activation, particularly in early phases of injury. In this context, the reduction of RhoA observedin our slice model following injury may reflect, at least in part early cytoskeletal disassembly and loss of axonal integrity rather than a contradiction of the canonical RhoA/ROCK activation paradigm **(Fig. 3E)**. Importantly, the SCS *in vitro* model captures early to subacute phases of SCI but lacks key components of the *in vivo* injury environment, including infiltrating immune cells, myelin debris, and sustained extracellular inhibitory signaling. As these factors are known to contribute to prolonged RhoA activation, their absence may result in altered or transient RhoA dynamics. Treatment with sEVs restored RhoA expression to control levels, in parallel with changes with downregulation of MAP2 and NF-H **(Fig. 3A, B)** and increased pSTAT3 **(Fig. 4E)** signaling, suggesting partial stabilization of cytoskeletal architecture and modulation of early injury responses. Treatment with iMR-90-sEVs resulted in a more pronounced decrease of MAP2 and NF-H **(Fig. 3A, B)** and an increased S100β response compared to SPC-01-sEVs. In contrast, SPC-01-sEVs led to a stronger elevation of RhoA **(Fig. 3E),** while the modulation of Nogo-A **(Fig. 3D)** showed a slight preference for iMR-90-sEVs, suggesting that iMR-90-sEVs might have a stronger effect on cytoskeletal modulation post-SCI.

Apoptosis is a major component of secondary injury following SCI, mediated through intrinsic and extrinsic signaling pathways. The extrinsic pathway is triggered by death receptor activation, whereas the intrinsic pathway is regulated by the balance between pro-apoptotic Bax and anti-apoptotic Bcl-xL, with Bax promoting cytochrome c release and Bcl-xL preserving mitochondrial integrity, stabilizing microtubules, and supporting myelin and oligodendrocyte survival (Galluzzi et al., 2009). Injury-induced Ca²⁺ influx activates proteolytic enzymes such as caspases and calpains, degrading cellular proteins and compromising cytoskeletal integrity, with caspase-3 as a central mediator of intrinsic pathway. It is rapidly activated in neurons and oligodendrocytes, leading to cytoskeletal degradation, DNA fragmentation, and disruption of cellular repair processes (Citron et al., 2000). MSC- (W. Liu et al., 2019) and NSC-sEVs (Rong et al., 2019) have been shown mitigate apoptosis in SCI, reducing Bax and cleaved caspase-3 while increasing Bcl-xL, promoting autophagy and neuronal survival. In our SCS model, untreated slices displayed a rapid increase in cleaved caspase-3 **(Fig. 4A)** and Bax **(Fig. 4B)** levels, accompanied by a decrease in Bcl-xL **(Fig. 4C)**, reflecting massive apoptosis. Treatment with SPC-01- and iMR-90-sEVs reduced cleaved caspase-3 and, to a greater extent, Bax. Bcl-xL levels remained largely unchanged in the iMR-90-sEV treated group compared to the SCI **(Fig. 4C)** but were markedly elevated in SPC-01-sEVs group, suggesting a stronger anti-apoptotic effect of SPC-01-sEVs.

The PI3K/Akt/mTOR pathway regulates cell growth, survival, and homeostasis and plays a key role in SCI, controlling inflammation, apoptosis, and immune cell activation during the acute phase, and supporting neural repair in subacute and chronic phases (Park et al., 2008; Sekiguchi et al., 2012). PTEN is a negative regulator of this pathway, dephosphorylating phosphatidylinositol-3,4,5-trisphosphate (PIP₃), thereby inhibiting Akt activation and downstream mTOR signaling (Georgescu, 2010). High PTEN levels post-SCI exacerbate cytoskeletal disruption and promote pro-apoptotic signaling via Bax and caspases. PTEN inhibition or deletion enhances PI3K/Akt signaling, reduces apoptosis, and promotes axonal regrowth and neural repair, highlighting its potential as a therapeutic target, although chronic-phase glial scar formation may limit regeneration (Danilov & Steward, 2015; Kath et al., 2018; C. L. Xiao et al., 2022; Yin et al., 2018).Previous studies in SCI models have shown that specific miRNAs enriched in extracellular vesicles can regulate PTEN expression and thereby modulate PI3K/Akt signaling, enhancing Bcl-xL expression, and suppressing Bax and caspase activity (Chen et al., 2021; He et al., 2020; Lv et al., 2024), ultimately promoting neuronal survival, neurite outgrowth, and cytoskeletal integrity (W. Y. Li et al., 2021) For example, exosome-associated miR-29b-3p (X. Xiao et al., 2021) and miR-26a (Chen et al., 2021) target PTEN and activate the Akt/mTOR pathway, contributing to improved neuronal survival and functional recovery. Similarly, miR-212-3p (Guan et al., 2021) miR-92a-3p (He et al., 2020) and miR-21 (Kang et al., 2019) have been shown to regulate PTEN-dependent PI3K/Akt signaling and reduce apoptosis following SCI. Together, these findings support PTEN as a recurrent target of miRNAs associated with extracellular vesicles, providing a plausible mechanistic link to the PTEN downregulation observed in our model. However, direct validation of specific miRNA–PTEN interactions was beyond the scope of the present study. Consistent with these observations, our results show that PTEN levels were slightly elevated in SCS following SCI induction **(Fig. 4D)**, correlating with increased Bax **(Fig. 4B)**, cleaved caspase-3 (**Fig. 4A**), and MAP2/NF-H disruption **(Fig. 3A, 3B)**. Vesicle treatment markedly reduced PTEN, decreased cleaved caspase-3, and shifted the Bax and Bcl-xL levels toward anti-apoptotic signaling, consistent with previous reports (X. G. Li et al., 2019). These PI3K/Akt-mediated pro-survival effects provide a foundation for subsequent activation of JAK/STAT3 pathway, a key regulator in SCI that coordinates inflammatory responses, cell survival, and post-injury repair. STAT3 activation promotes transcription of anti-apoptotic genes, including Bcl-xL, supporting neuronal survival, and limiting apoptosis. In astrocytes, STAT3 phosphorylation drives formation of a structured glial scar, restricting inflammatory infiltration and lesion expansion, contributing to functional recovery (Okada et al., 2006). Conversely, deletion of STAT3 in astrocytes disrupts scar organization and exacerbates neuronal loss (Wanner et al., 2013), highlighting its dual role: protective in the acute phase but potentially inhibitory in the chronic phase. The JAK2/STAT3 pathway is activated rapidly after SCI, peaking within ∼12 hours and declining by 24 hours, indicating a crucial role in early regulation of apoptosis, autophagy, and inflammatory signaling (Abbaszadeh et al., 2020). In our SCI model, pSTAT3 levels remained near control levels 3 days post-injury, reflecting the decline of early STAT3 activation following the acute phase **(Fig. 4E)**, or, similarly to downregulated RhoA post-SCI **(Fig. 3E)**, early cytoskeletal disassembly, and loss of axonal integrity. sEVs treatment strongly upregulated pSTAT3 levels, coinciding with elevated Bcl-xL **(Fig. 4C)** and cytoskeletal stabilization. This coordinated activation of PI3K/Akt via PTEN inhibition and enhanced pSTAT3 phosphorylation is consistent with a putative synergistic interaction, through which sEVs may promote neuroprotection, support cytoskeletal stability, and promote astrocytic responses during the early phases of SCI.

Overall, iMR-90-sEVs appeared to exert a stronger effect on cytoskeletal proteins than SPC-01-sEVs **(Fig. 3)**, whereas SPC-01-sEVs exhibited a more pronounced anti-apoptotic effect, reflected by higher Bcl-xL levels **(Fig. 4C)** and a slightly greater reduction in cleaved caspase-3 and Bax compared to the SCI group **(Fig. 4A, 4B)**. The effects on pSTAT3 and PTEN were largely similar, with PTEN significantly downregulated and pSTAT3 significantly upregulated compared to the SCI group **(Fig. 4D, E)**. These observations suggest differences in the effects of SPC-01-sEVs and iMR-90-sEVs, which cannot be fully elucidated without a comprehensive analysis of the miRNA cargo of these vesicles.

Collectively, our findings indicate that sEV treatment induces a coordinated neuroprotective response, with early reactive gliosis likely reflecting containment of the lesion rather than mature scar formation. Vesicles increased STAT3 phosphorylation, reduced PTEN, and elevated Bcl-xL, supporting survival signaling via PI3K/Akt, and attenuating apoptosis, as reflected by decreased cleaved caspase-3 and Bax. Normalization of MAP2 and NF-H, along recovery of RhoA and reduction of Nogo-A, indicates modulation of cytoskeletal organization and structural remodeling. Importantly, given the SCS model lacks systemic immune cell infiltration, these effects are likely mediated by direct interactions with resident neurons and glia. Although limited to a 72-hour time point and an *in vitro* system that does not fully recapitulate the in vivo inflammatory milieu, the model captures early intrinsic injury responses. Together, these findings suggest that early sEV treatment during the acute SCI may promote neuronal survival, cytoskeletal stabilization, and activation of early neuroprotective signaling pathways within injured spinal cord tissue.

Early changes in PTEN, STAT3 phosphorylation, Bcl-xL, Nogo-A, and RhoA are considered mechanistic indicators of whether the post-injury environment is shifting toward repair rather than degeneration, and therefore can provide predictive information about long-term functional recovery after SCI. Reduced PTEN suggests activation of PI3K/Akt signaling, which promotes neuronal survival and axonal growth; sustained suppression of PTEN has been associated with enhanced regenerative capacity and improved functional outcomes in SCI models (Cho et al., 2020; Ohtake et al., 2015). Increased STAT3 phosphorylation reflects activation of a context-dependent repair pathway that can support neuronal survival and modulate inflammatory responses, contributing to a more permissive regenerative environment when appropriately regulated (R. Liu et al., 2023). Elevated Bcl-xL indicates reduced apoptosis, which is critical in the acute phase because preservation of neurons and oligodendrocytes directly determines the substrate available for later circuit reorganization and conduction recovery (Cittelly et al., 2007). In parallel, decreased Nogo-A and normalized RhoA suggest attenuation of inhibitory myelin-associated signaling and cytoskeletal constraints that normally prevent axon extension (Waliullah et al., 2025). This shift reduces growth cone collapse and facilitates axonal sprouting and structural plasticity across and around the lesion site.

Together, these molecular changes indicate that sEV treatment promotes a coordinated pro-survival and pro-growth state. If maintained over time, such a state increases the likelihood of white matter preservation, enhances axonal regrowth, and improved synaptic connectivity. These structural effects are the biological basis for later improvements in motor coordination and locomotor function. Thus, early modulation of PTEN/STAT3/Bcl-xL alongside reduced Nogo-A/RhoA is not only a marker of acute neuroprotection but also a mechanistically grounded predictor of potential long-term functional recovery after spinal cord injury.

A limitation of this study is the incomplete characterization of the rat spinal cord slice (SCS) *in vitro* SCI model. Due to technical constraints and time limitations, a full longitudinal and multi-layered characterization could not be performed, and some aspects of cellular dynamics and injury evolution were not exhaustively assessed, particularly at later time points and in terms of broader functional network changes. Although the SCS model preserves key features of native spinal cord cytoarchitecture and enables controlled investigation of early post-injury responses, it does not fully reproduce the complexity of *in vivo* SCI. In particular, the absence of systemic immune cell infiltration and circulating immune interactions is a significant limitation, reflected in the progressive decline of microglial signal over time. In addition, the experimental design primarily captured late acute to early subacute molecular events (3 days post-injury) and therefore did not address chronic processes such as long-term axonal remodeling, glial scar maturation, or functional recovery.

Accordingly, while the model is suitable for studying early molecular and cellular responses to SCI and the effects of sEVs on neuronal survival and cytoskeletal regulation, it does not allow direct conclusions regarding long-term functional outcomes. Future studies with extended time courses and more comprehensive phenotyping would be needed to fully characterize the temporal evolution of injury and therapeutic responses

## 5 Conclusion

Spinal cord injury remains a major challenge in neuroregenerative medicine, as current therapies provide only limited support for neural repair and regeneration. Here, we characterized small extracellular vesicles derived from neural stem cells (SPC-01 and iMR-90-derived neural precursors) and evaluated their neuroprotective and anti-apoptotic effects using rat organotypic spinal cord slices as an *in vitro* SCI model. Treatment with sEVs modulated PTEN/STAT3-associated survival pathways, reduced apoptotic signaling, and contributed to stabilization of neuronal cytoskeletal proteins and reduction of inhibitory Nogo-A signaling. These findings indicate that neural stem cell-derived sEVs can promote early neuroprotective responses in injured spinal cord tissue, highlighting their potential as a therapeutic strategy for SCI.

## 7 Ethics statement

The experimental procedures were approved by the Ethics Committee of the IEM ASCR and are in accordance with the Directive of the European Commission of November 24, 1986 (86/609/EEC) on the use of animals in research

## Acknowledgments

We acknowledge CF Biophysic of CIISB, Instruct-CZ Centre, supported by MEYS CR (LM2023042) and European Regional Development Fund-Project „Innovation of Czech Infrastructure for Integrative Structural Biology“(No. CZ.02.01.01/00/23_015/0008175). We would also like to acknowledge Imaging Methods Core Facility at BIOCEV.

## 9 Data Availability Statement

The original contributions presented in the study are included in the article/Supplementary material. The cell line iPSC-NP (iMR-90) present in this study was obtained from Brigitte Onteniente, who provided it within the European project STEMS. Recently, the cell line is available as PCi-NPC from company Phenocell (Grasse, France). The cell line SPC-01 was obtained from Prof. Jack Price, who provided it within the European project RESCUE. Further inquiries can be directed at the corresponding authors.

---

**Supplementary table 1:** List of primary antibodies used for Western Blot and Immunohistochemistry (IHC). All antibodies are from Cell Signaling Technology, Danvers, MA, USA, unless stated otherwise.

| Antibody | IHC | Western blot | Molecular weight (kDa) | Catalog number |
| --- | --- | --- | --- | --- |
| **MAP2** | 1:50 | 1:1000, BSA | 75, 82, 280 | 4542 |
| **Neurofilament-H** | 1:400 | 1:1000, milk | 180-220 | 2836 |
| **Nogo-A** | - | 1:1000, BSA | 180 | 13401 |
| **Vinculin** | - | 1:1000, BSA | 124 | 13901 |
| **Alix** | - | 1:1000, milk | 90-100 | 92880 |
| **Calnexin** | - | 1:1000, BSA | 90 | 10427-2-AP<br>(proteintech) |
| **pSTA3** | - | 1:1000, BSA | 79, 86 | 9145 |
| **STAT3** | - | 1:1000, milk | 79, 86 | 9139 |
| **c-myc** | - | 1:1000, BSA | 57-65 | 18583 |
| **Vimentin** | 1:200 | - | 57 | 5741 |
| **βIII-tubulin** | 1:400 | - | 55 | 5568 |
| **PTEN** | - | 1:1000, BSA | 54 | 9559 |
| **GFAP** | - | 1:1000, milk | 50 | 12389 |
| **GFAP** | 1:200 | - | 50 | C9205<br>(Sigma-Aldrich) |
| **TSG101** | - | 1:1000, BSA | 49 | 77452<br>(Novus Biologicals) |
| **β-actin** | - | 1:2000, TBST | 42 | A2228<br>(Sigma-Aldrich) |
| **Annexin V** | - | 1:1000, BSA | 30 | 8555 |
| **Bcl-xL** | - | 1:1000, milk | 30 | 2764 |
| **CD9** | - | 1:1000, BSA | 22, 24, 35 | 13403 |
| **CD81** | - | 1:1000, BSA | 8, 25 | 65805<br>(Novus Biologicals) |
| **CD63** | - | 1:1000, BSA | 6, 25 | 42225<br>(Novus Biologicals) |
| **RhoA** | - | 1:1000, BSA | 21 | 2117 |
| **Bax** | - | 1:1000, BSA | 20 | 14796 |
| **Cleaved Caspase-3** | - | 1:1000, BSA | 17, 19 | 9664 |
| **Iba1** | 1:200 | - | 17 | 17198 |
| **S100β** | 1:400 | 1:1000, BSA | 10 | 90393 |

**Supplementary table 2:**
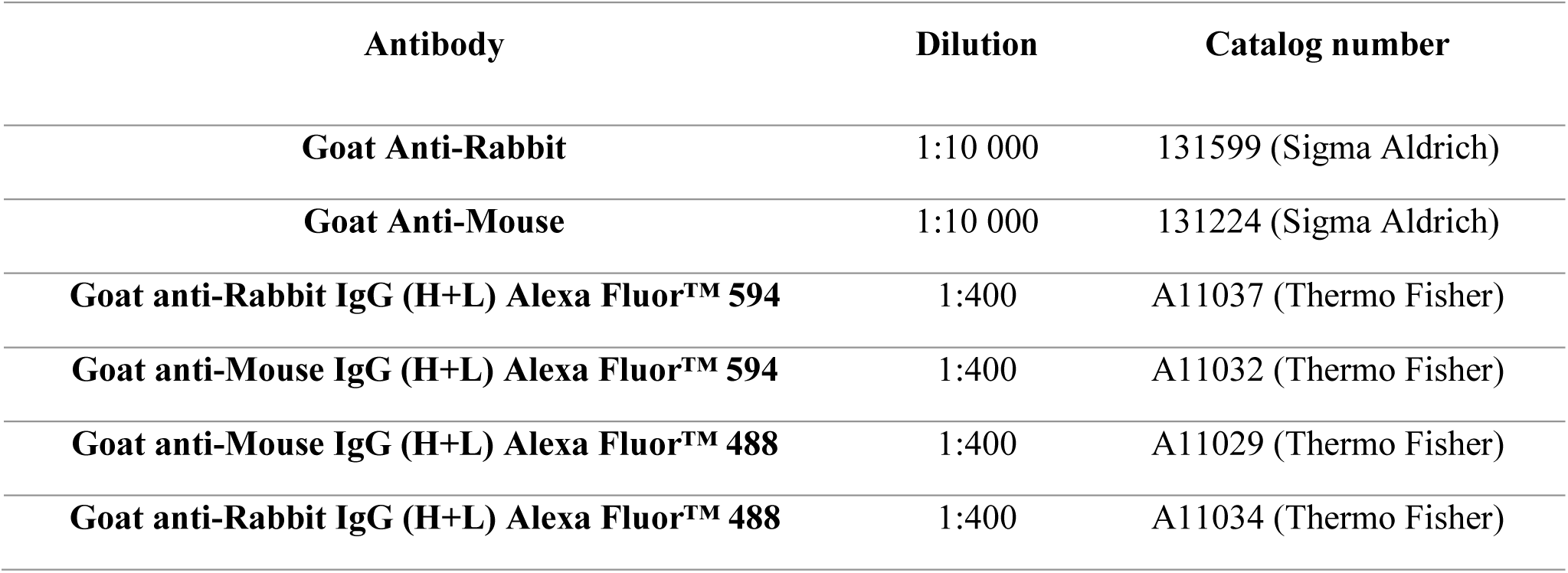
List of secondary antibodies used for Western Blot and Immunohistochemistry (IHC).

