## Supplementary File 1 for "Neural stem cell–derived extracellular vesicles drive early neuroprotective and anti-apoptotic responses in spinal cord injury organotypic slices"

### Slide 1
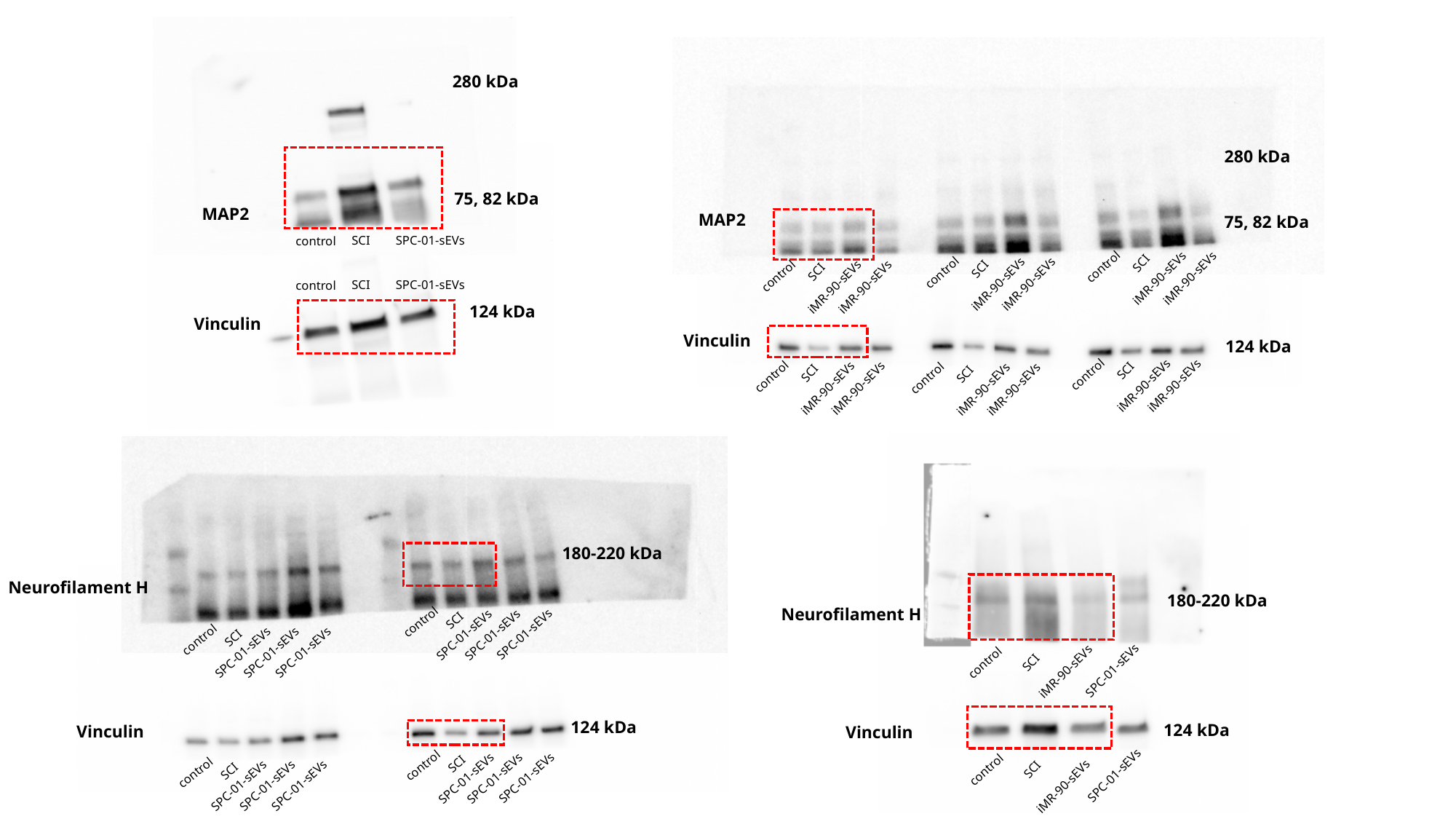

280 kDa
MAP2
SCI
SPC-01-sEVs
control
75, 82 kDa
280 kDa
MAP2
75, 82 kDa
iMR-90-sEVs
SCI
control
iMR-90-sEVs
iMR-90-sEVs
SCI
control
iMR-90-sEVs
iMR-90-sEVs
SCI
control
iMR-90-sEVs
SCI
SPC-01-sEVs
control
Vinculin
124 kDa
iMR-90-sEVs
SCI
control
iMR-90-sEVs
iMR-90-sEVs
SCI
control
iMR-90-sEVs
Vinculin
iMR-90-sEVs
SCI
control
iMR-90-sEVs
124 kDa
180-220 kDa
Neurofilament H
SCI
control
iMR-90-sEVs
180-220 kDa
Neurofilament H
SCI
control
SPC-01-sEVs
SPC-01-sEVs
SPC-01-sEVs
SCI
control
SPC-01-sEVs
SPC-01-sEVs
SPC-01-sEVs
124 kDa
Vinculin
SCI
control
iMR-90-sEVs
124 kDa
Vinculin
SCI
control
SCI
SPC-01-sEVs
control
SPC-01-sEVs
SPC-01-sEVs
SPC-01-sEVs
SPC-01-sEVs
SPC-01-sEVs
SPC-01-sEVs
SPC-01-sEVs

### Slide 2
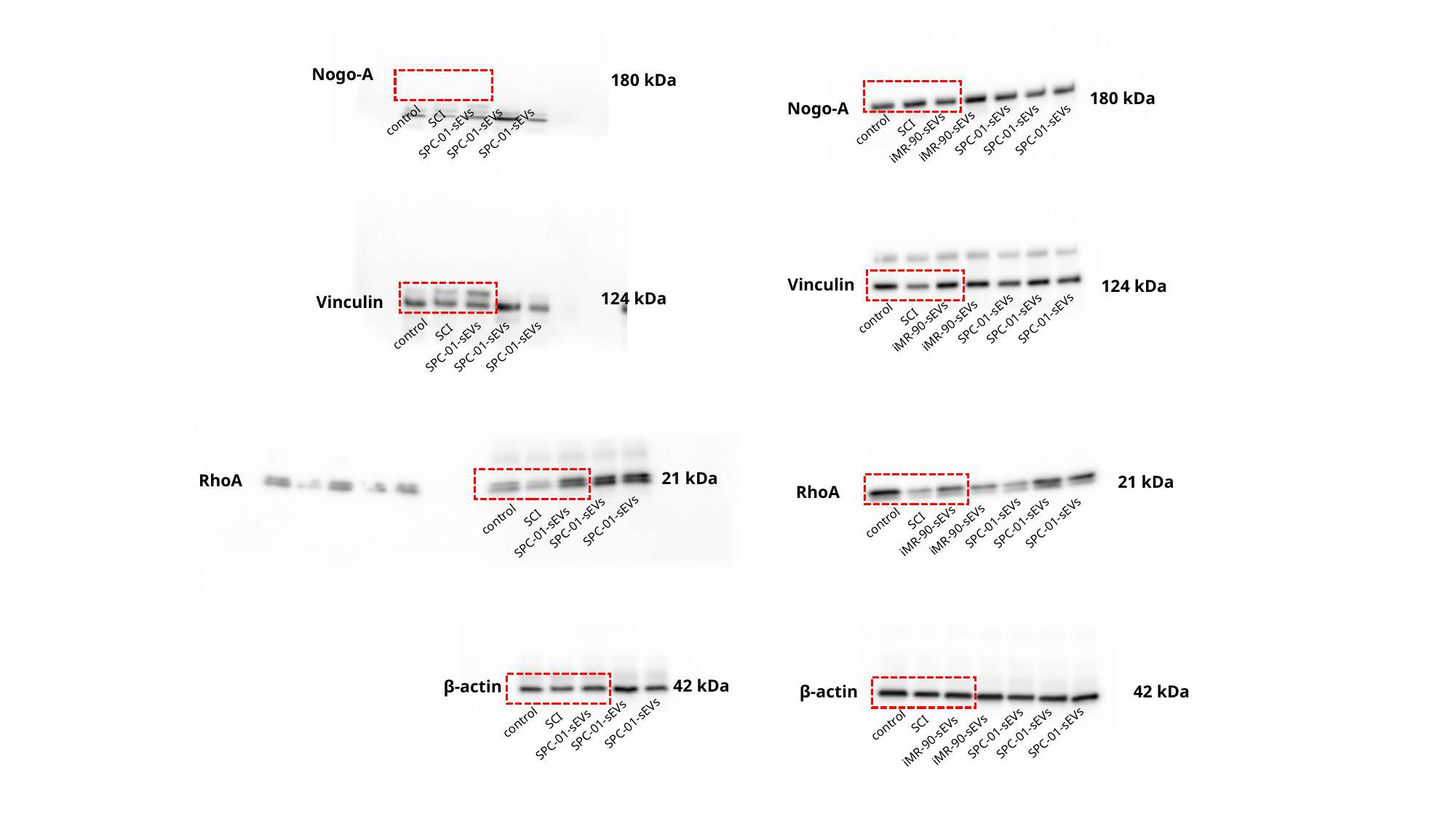

180 kDa
Nogo-A
SPC-01-sEVs
SPC-01-sEVs
SPC-01-sEVs
SCI
control
iMR-90-sEVs
iMR-90-sEVs
Nogo-A
180 kDa
SCI
control
SPC-01-sEVs
SPC-01-sEVs
SPC-01-sEVs
124 kDa
Vinculin
SCI
control
SPC-01-sEVs
SPC-01-sEVs
SPC-01-sEVs
Vinculin
124 kDa
SPC-01-sEVs
SPC-01-sEVs
SPC-01-sEVs
SCI
control
iMR-90-sEVs
iMR-90-sEVs
21 kDa
RhoA
SCI
SPC-01-sEVs
SPC-01-sEVs
control
SPC-01-sEVs
21 kDa
RhoA
SPC-01-sEVs
SPC-01-sEVs
SPC-01-sEVs
SCI
control
iMR-90-sEVs
iMR-90-sEVs
42 kDa
β-actin
SCI
SPC-01-sEVs
SPC-01-sEVs
control
SPC-01-sEVs
β-actin
42 kDa
SCI
control
SPC-01-sEVs
SPC-01-sEVs
SPC-01-sEVs
iMR-90-sEVs
iMR-90-sEVs

### Slide 3
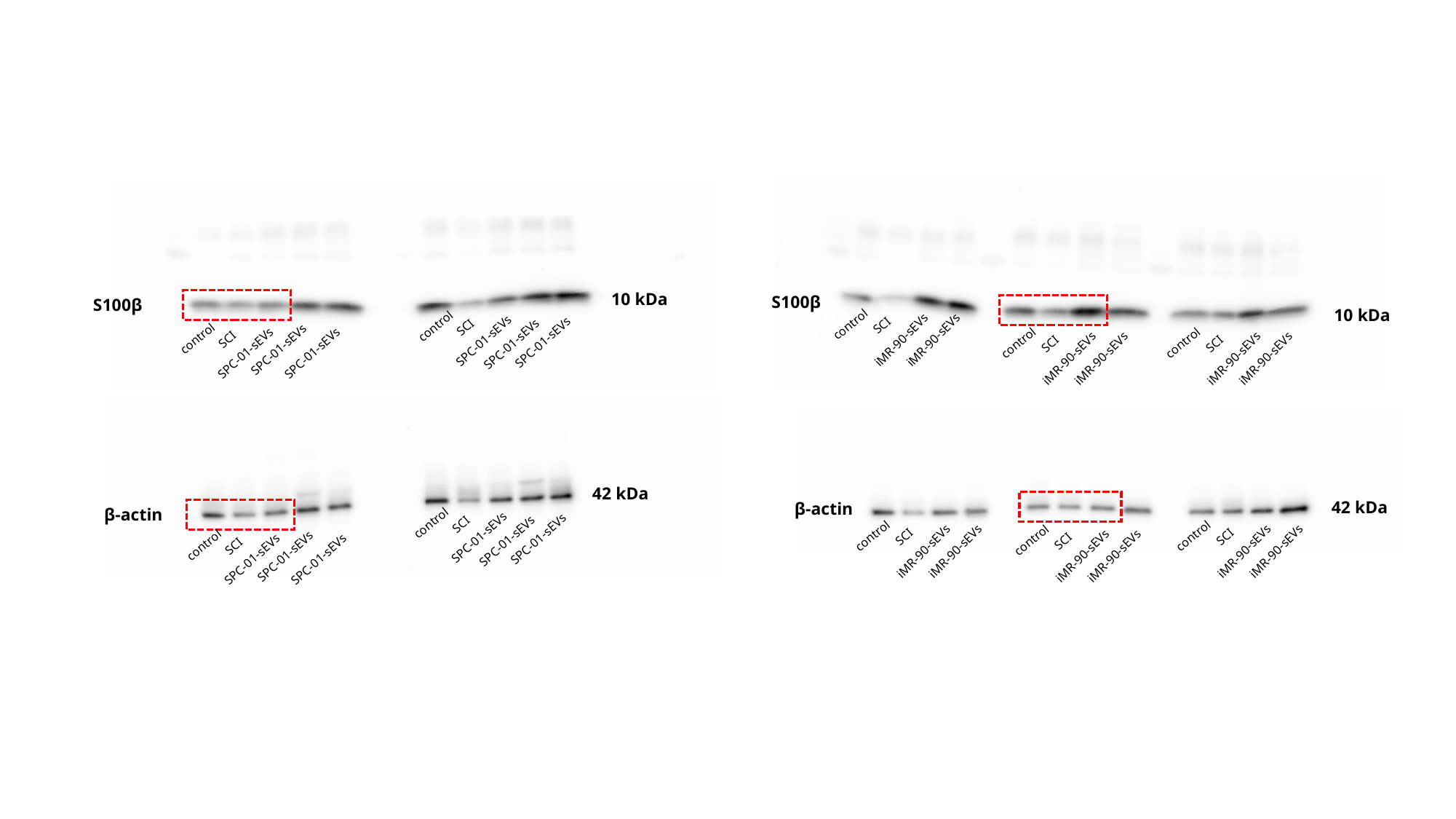

S100β
10 kDa
SCI
control
iMR-90-sEVs
iMR-90-sEVs
SCI
SCI
control
control
iMR-90-sEVs
iMR-90-sEVs
iMR-90-sEVs
iMR-90-sEVs
10 kDa
S100β
SCI
control
SPC-01-sEVs
SPC-01-sEVs
SPC-01-sEVs
SCI
control
SPC-01-sEVs
SPC-01-sEVs
SPC-01-sEVs
42 kDa
β-actin
SCI
control
SPC-01-sEVs
SPC-01-sEVs
SPC-01-sEVs
SCI
control
SPC-01-sEVs
SPC-01-sEVs
SPC-01-sEVs
42 kDa
β-actin
SCI
SCI
control
control
SCI
control
iMR-90-sEVs
iMR-90-sEVs
iMR-90-sEVs
iMR-90-sEVs
iMR-90-sEVs
iMR-90-sEVs

### Slide 4
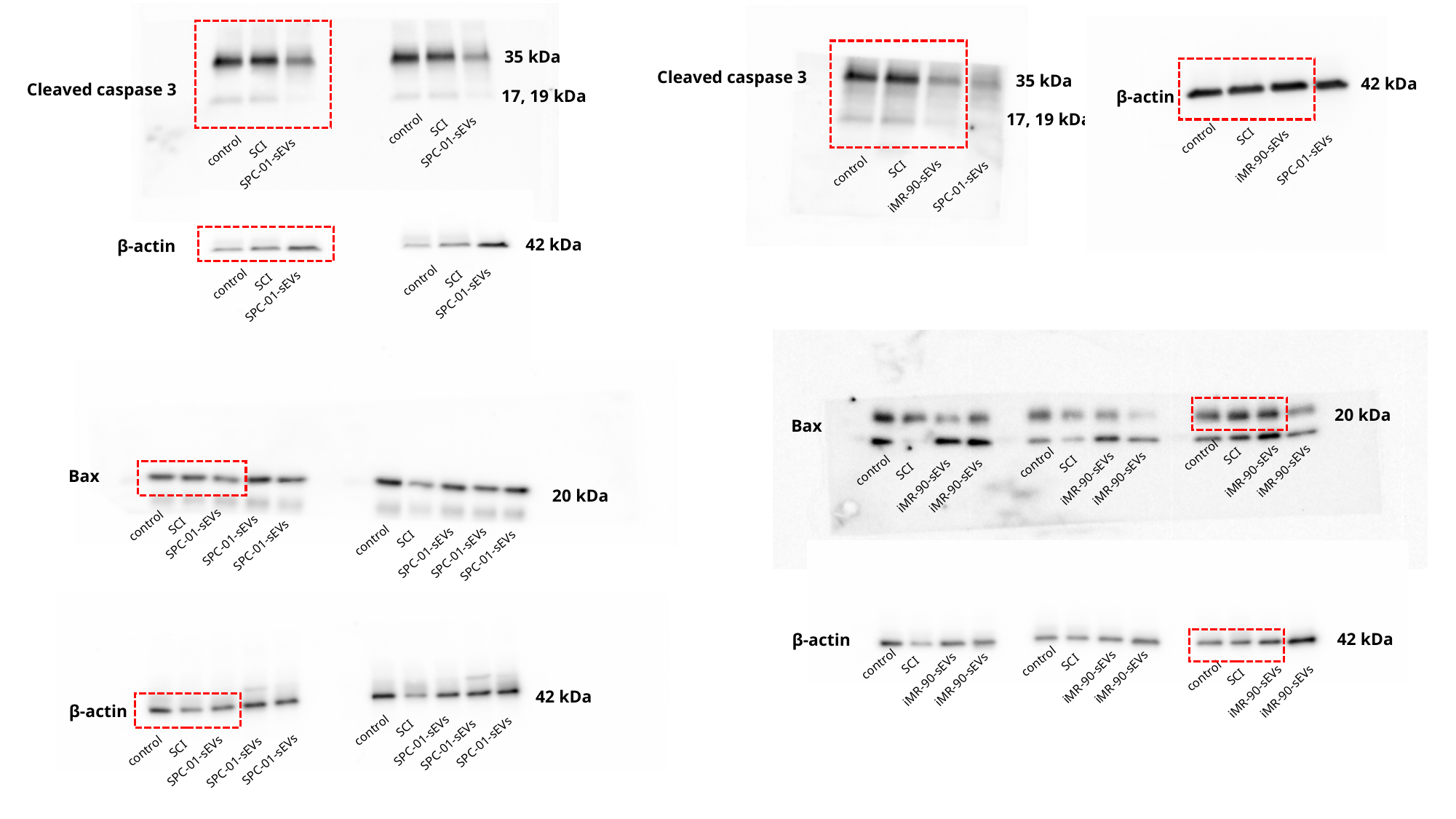

35 kDa
Cleaved caspase 3
17, 19 kDa
SCI
control
SPC-01-sEVs
SCI
control
SPC-01-sEVs
Cleaved caspase 3
35 kDa
17, 19 kDa
SCI
control
SPC-01-sEVs
iMR-90-sEVs
42 kDa
β-actin
SCI
control
SPC-01-sEVs
iMR-90-sEVs
42 kDa
β-actin
SCI
control
SPC-01-sEVs
SCI
control
SPC-01-sEVs
20 kDa
Bax
SCI
control
SCI
control
iMR-90-sEVs
iMR-90-sEVs
SCI
control
iMR-90-sEVs
iMR-90-sEVs
iMR-90-sEVs
iMR-90-sEVs
Bax
20 kDa
SCI
control
SPC-01-sEVs
SCI
SPC-01-sEVs
SPC-01-sEVs
control
SPC-01-sEVs
SPC-01-sEVs
SPC-01-sEVs
42 kDa
β-actin
SCI
control
SCI
control
iMR-90-sEVs
iMR-90-sEVs
iMR-90-sEVs
iMR-90-sEVs
SCI
control
iMR-90-sEVs
iMR-90-sEVs
42 kDa
β-actin
SCI
control
SPC-01-sEVs
SPC-01-sEVs
SPC-01-sEVs
SCI
control
SPC-01-sEVs
SPC-01-sEVs
SPC-01-sEVs

### Slide 5
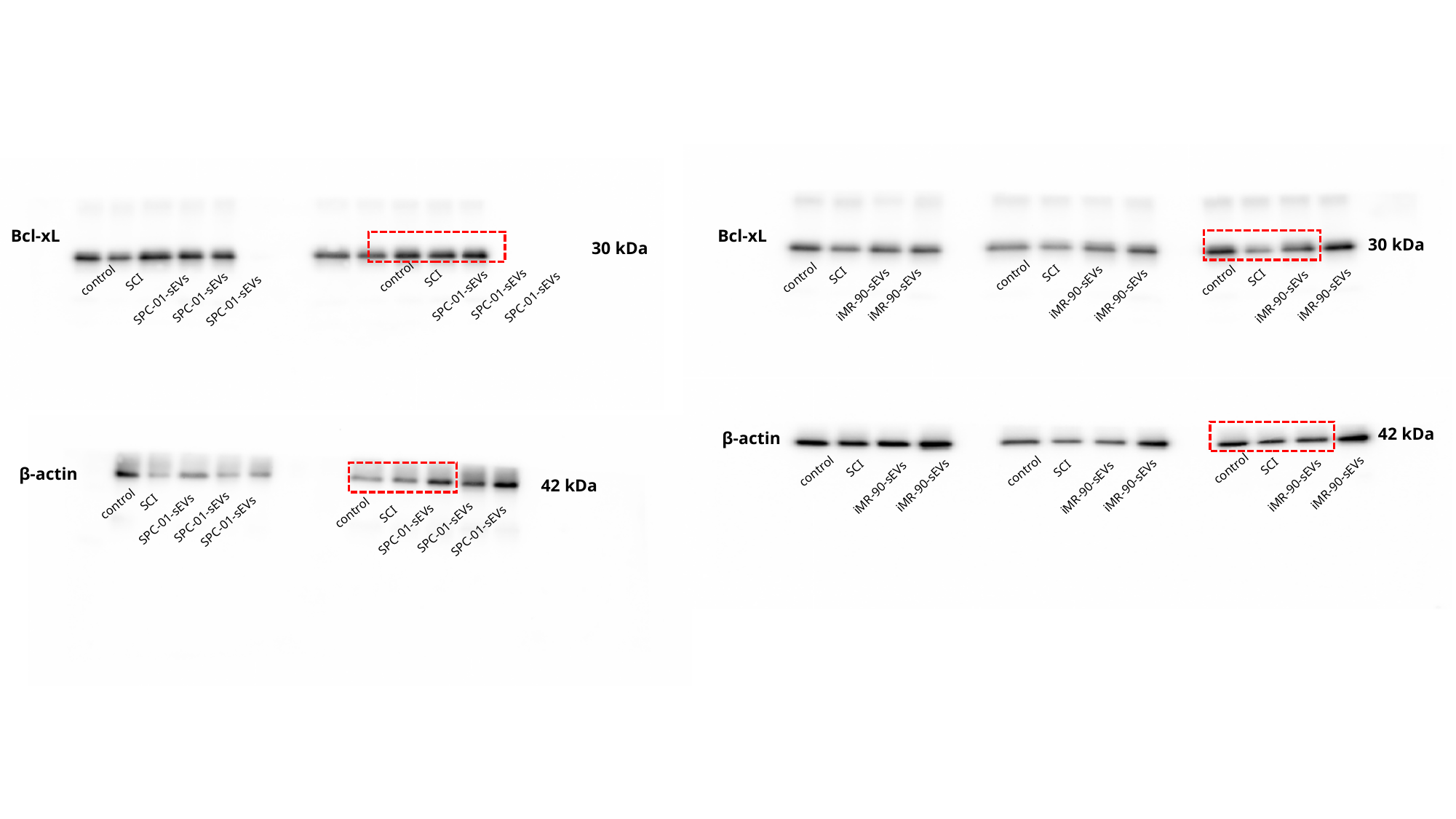

Bcl-xL
30 kDa
SCI
SCI
SCI
control
control
control
iMR-90-sEVs
iMR-90-sEVs
iMR-90-sEVs
iMR-90-sEVs
iMR-90-sEVs
iMR-90-sEVs
Bcl-xL
30 kDa
SCI
control
SCI
control
SPC-01-sEVs
SPC-01-sEVs
SPC-01-sEVs
SPC-01-sEVs
SPC-01-sEVs
SPC-01-sEVs
42 kDa
β-actin
SCI
SCI
SCI
control
control
control
iMR-90-sEVs
iMR-90-sEVs
iMR-90-sEVs
iMR-90-sEVs
iMR-90-sEVs
iMR-90-sEVs
β-actin
42 kDa
SCI
control
SCI
SPC-01-sEVs
control
SPC-01-sEVs
SPC-01-sEVs
SPC-01-sEVs
SPC-01-sEVs
SPC-01-sEVs

### Slide 6
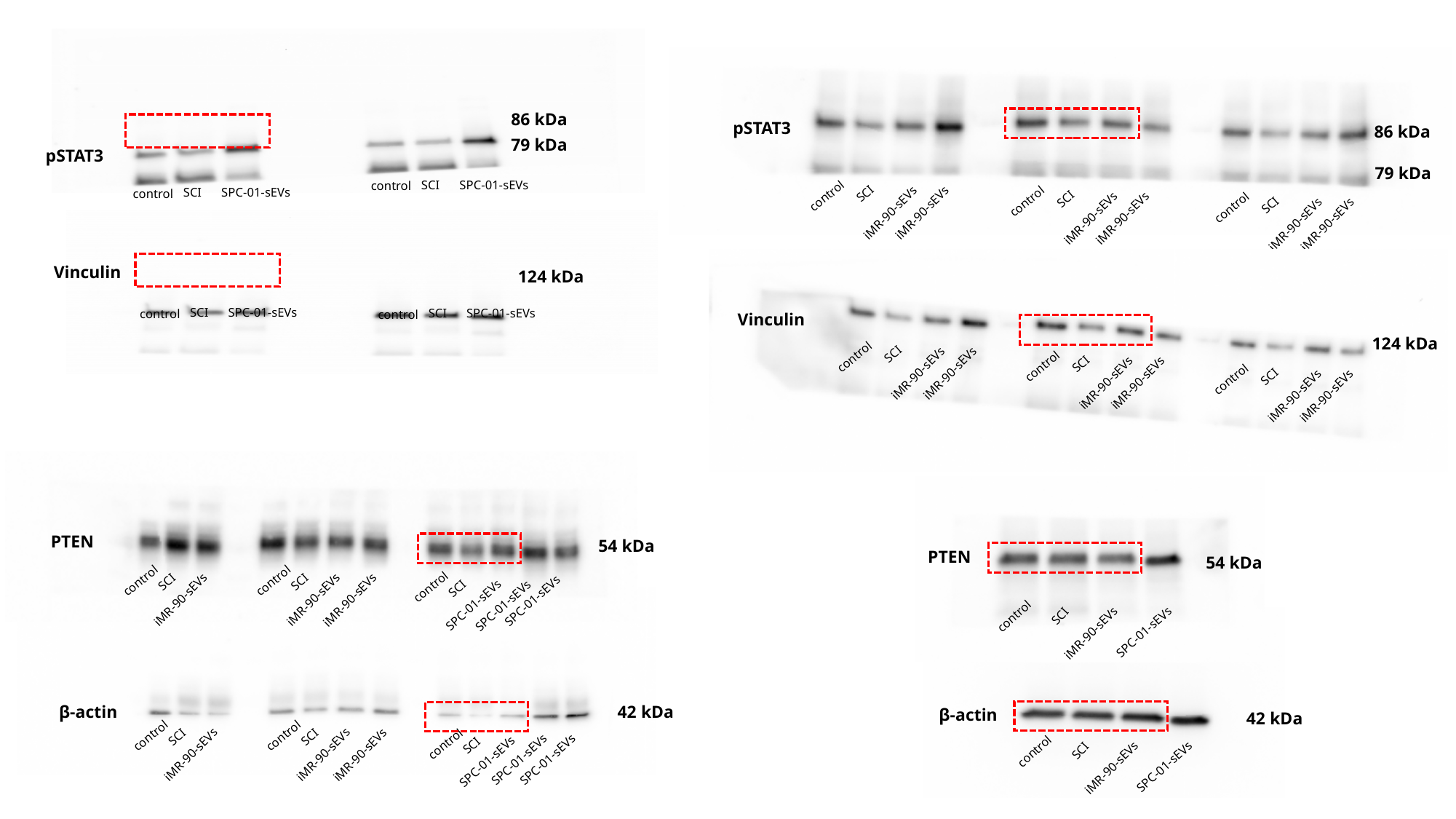

86 kDa
79 kDa
pSTAT3
SCI
SPC-01-sEVs
control
SCI
SPC-01-sEVs
control
pSTAT3
86 kDa
79 kDa
SCI
SCI
control
control
SCI
iMR-90-sEVs
iMR-90-sEVs
control
iMR-90-sEVs
iMR-90-sEVs
iMR-90-sEVs
iMR-90-sEVs
Vinculin
124 kDa
SCI
SPC-01-sEVs
SCI
SPC-01-sEVs
control
control
Vinculin
124 kDa
SCI
control
SCI
control
iMR-90-sEVs
iMR-90-sEVs
SCI
iMR-90-sEVs
iMR-90-sEVs
control
iMR-90-sEVs
iMR-90-sEVs
PTEN
54 kDa
SCI
SCI
control
control
SCI
control
iMR-90-sEVs
iMR-90-sEVs
iMR-90-sEVs
SPC-01-sEVs
SPC-01-sEVs
SPC-01-sEVs
PTEN
SCI
control
iMR-90-sEVs
SPC-01-sEVs
54 kDa
42 kDa
β-actin
SCI
SCI
control
control
SCI
control
iMR-90-sEVs
iMR-90-sEVs
iMR-90-sEVs
SPC-01-sEVs
SPC-01-sEVs
SPC-01-sEVs
β-actin
42 kDa
SCI
control
iMR-90-sEVs
SPC-01-sEVs
